# Bacterial fitness for plant colonization is influenced by plant growth substrate

**DOI:** 10.1101/2025.01.16.633496

**Authors:** Marta Torres, Morgan N. Price, Albina Kashanova, Suzanne M. Kosina, Kateryna Zhalnina, Trent R. Northen, Adam M. Deutschbauer

## Abstract

- Bacteria colonize plants and contribute to their health. Despite advances in our understanding of bacterial plant colonization, the extent to which growth substrate influences the molecular mechanisms enabling bacteria to efficiently colonize plants remains poorly understood.
- To evaluate if genes important for plant colonization are influenced by substrate, we used randomly barcoded transposon mutagenesis sequencing (RB-TnSeq) in *Paraburkholderia graminis* OAS925, an efficient rhizosphere colonizer, and *Brachypodium distachyon* plants grown in six different substrates.
- Of the 382 rhizosphere colonization genes that we identified in strain OAS925, 348 genes (91.1%) are dependent on the growth substrate evaluated, and 34 genes (8.9%) are shared across all the substrates. To further characterize the plant colonization fitness genes, we compared our data to growth in 110 distinct *in vitro* conditions. To validate our results, we used individual mutants of selected genes and tested them in different growth substrates. We also show the influence of substrate on colonization by *P. phytofirmans* PsJN, *Variovorax* sp. OAS795 and *Rhizobium* sp. OAE497.
- Our data confirms that bacterial fitness for plant colonization is strongly influenced by plant growth substrate type, and highlights the importance of taking this parameter into consideration when engineering bacterial strains for improved host colonization.

## 1. Introduction

Plants release 20-40% of photosynthesis-derived carbon into the rhizosphere, the thin soil layer surrounding the roots (Kuzyakov & Domanski, 2000; Jones *et al*., 2009). These root-derived compounds include a wide variety of substances that originate from dying root cap border cells, mucilages, volatiles and root exudates, among others (Dennis *et al*., 2010). Root exudates typically comprise primary metabolites such as sugars, amino acids, organic acids, fatty acids, inorganic ions, purines, and secondary metabolites (Dakora & Phillips, 2002a; Bais *et al*., 2006; Baetz & Martinoia, 2014). These can act as signaling molecules to attract beneficial organisms and to repel pathogens. They can also be used as nutrients by soil microorganisms, which translates into a faster proliferation of microbial cells in the rhizosphere compared to non-rhizosphere soil. This influence on the microorganisms in the immediate vicinity of the root has been referred to as the ‘rhizosphere effect’ (Berendsen *et al*., 2012; Pascale *et al*., 2020). In fact, root-derived nutrients mediate plant-plant and plant-microbe interactions and are one of the main drivers structuring rhizosphere microbial communities along with host plant immunity (Bais *et al*., 2006; Haichar *et al*., 2008; Badri & Vivanco, 2009; Williams & Vries, 2020).

Among the factors that influence root metabolite composition (i.e. plant species, plant growth stage, environmental conditions, and root location), some of them interconnected, plant growth substrate is one of the least characterized (Yang & Crowley, 2000; Gransee & Wittenmayer, 2000; Hertenberger *et al*., 2002; van Dam & Bouwmeester, 2016; Zhalnina *et al*., 2018; Williams & Vries, 2020; McLaughlin *et al*., 2023). In a recent study, authors showed that soil chemistry has an impact on root morphology and exudate composition in *Brachypodium distachyon* plants (Neumann *et al*., 2014; Sasse *et al*., 2019). Interestingly, plant-derived substances can, in turn, modify the physical and chemical environment of the rhizosphere (Dakora & Phillips, 2002b; Oleghe *et al*., 2017; Naveed *et al*., 2018). Type of soil can also affect diffusion and adsorption of root-exuded compounds, therefore impacting nutrient availability for soil microorganisms (Ndzana *et al*., 2022), and microbial behavior, for instance influencing biofilm formation (Ma *et al*., 2017) and motility (Yang & Van Elsas, 2018). Taken together, it seems evident that the growth substrate should be considered when assessing the performance of beneficial bacteria on plants, or when conducting plant-microbe interaction studies.

The diverse community of soil microorganisms inhabiting the rhizosphere plays an important role in plant nutrition and protection against pathogens (Müller *et al*., 2016). Consequently, promoting plant growth by harnessing the soil microbiome stands as a promising and sustainable alternative to agrochemicals (Compant *et al*., 2024). In order to increase crop yield and quality, numerous efforts have aimed at both isolating beneficial members of the soil microbiota and better understanding how plant colonization by soil bacteria is shaped (Haichar *et al*., 2008; Schreiter *et al*., 2014; Torres *et al*., 2022; Tang *et al*., 2022). One needed step is having a genome-wide map of fitness determinants involved in plant colonization by a given bacterium (Cole *et al*., 2017), which offers a starting point for future actions such as the targeted improvement of plant colonization capabilities of beneficial bacteria, or the development of protection strategies against plant pathogens (Cole *et al*., 2017; Torres *et al*., 2022; Compant *et al*., 2024). Individual strains and microbial consortia (synthetic communities, SynComs) with promising biotechnological applications are typically selected based on laboratory assays conducted in defined plant media. Transferring lab research to the natural environment remains a challenge due to inconsistent field efficacy, since efficient plant colonization is context-dependent and can vary due to factors such as plant growth substrate (Elsas *et al*., 1986; Jindo *et al*., 2020; Li *et al*., 2022). This could be due to a systematic evaluation of the colonization and/or plant-growth stimulation abilities of beneficial bacteria in a non-natural type of substrate, often an agar-based gnotobiotic system (Cole *et al*., 2017; Vannier *et al*., 2023).

High-throughput techniques, such as randomly barcoded transposon mutagenesis sequencing (RB-TnSeq) (Wetmore *et al*., 2015) that combines the advantages of TnSeq (Opijnen *et al*., 2009) and rapid quantification of each transposon mutant using unique DNA barcodes, have been used to identify the fitness genetic determinants that are involved in a function of interest. In the last years, extensive studies using TnSeq and RB-TnSeq approaches have allowed the identification of genes involved in root and rhizosphere colonization in plant-associated species such as *Agrobacterium tumefaciens, Aquitalea magnusonii, Azoarcus olearius*, *Dickeya solani*, *Herbaspirillum seropedicae, Pseudomonas fluorescens, P. simiae*, *P. syringae*, *Rhizobium leguminosarum, Sinorhizobium meliloti*, and *Xanthomonas campestris* (Cole *et al*., 2017; Liu *et al*., 2018a; Helmann *et al*., 2019; Wheatley *et al*., 2020; Flores-Tinoco *et al*., 2020; Amaral *et al*., 2020; Torres *et al*., 2022; Ishizawa *et al*., 2022; Luneau *et al*., 2022; Robic *et al*., 2023). These studies were conducted in plants grown in a diversity of substrates, including plant growth media like Murashige & Skoog or Hoagland solution (Cole *et al*., 2017; Wheatley *et al*., 2020; Amaral *et al*., 2020; Ishizawa *et al*., 2022), perlite (Flores-Tinoco *et al*., 2020), potting mix (Torres *et al*., 2022; Robic *et al*., 2023), or mixes of clay-vermiculite (Amaral *et al*., 2020) or clay-perlite (Liu *et al*., 2018a). Globally, around 1-8% of the genome of these bacterial species was revealed to be involved in plant colonization. The identified genes belong to categories such as amino acid transport and metabolism, cell wall/membrane biogenesis and cell motility (Fabian *et al*., 2020; Torres *et al*., 2024), and they offer a promising starting point for targeted improvement of the colonization capabilities of plant-beneficial microbes (Cole *et al*., 2017) or for deterring soil pathogenic bacteria (Torres *et al*., 2022). Despite the valuable knowledge on bacterial plant colonization to which these studies have contributed, the extent to which the plant growth substrate used influences the outcome of studies on bacterial fitness for plant colonization has not been evaluated, hindering the comparison of plant colonization genes across organisms.

To determine how different plant growth substrates impact bacterial plant colonization fitness determinants, we used the plant *B. distachyon* and an RB-TnSeq library of the bacterium *Paraburkholderia graminis* OAS925 (henceforth OAS925). *B. distachyon* is a monocot used as a model for the study of a large number of plants with broad range of uses, such as emerging bioenergy crops (e.g. switchgrass, Chinese silver grass) or food crops (e.g. rice, wheat, barley) (Draper *et al*., 2001; Brkljacic *et al*., 2011). OAS925 is an isolate from switchgrass fields (Coker *et al*., 2022) that has been shown to dominate the population of *B. distachyon* roots and rhizosphere of plants inoculated with a 17-member SynCom (Lin *et al*., 2023; Novak *et al*., 2024). *P. graminis* is positioned in the cluster of plant-beneficial *Burkholderia/Paraburkholderia* species (Estrada-de los Santos *et al*., 2013; Sawana *et al*., 2014). Members of the order Burkholderiales, and more precisely from the *Burkholderiaceae* family, are some of the most common inhabitants of the rhizosphere (Compant *et al*., 2010; Weisskopf *et al*., 2011; Li *et al*., 2014). They can use numerous plant-derived compounds as nutrient sources (Estrada-De Los Santos *et al*., 2001; Kost *et al*., 2014; Korenblum *et al*., 2020), tolerate the acidic pH encountered in rhizosphere (Stopnisek *et al*., 2014; Ma *et al*., 2022), and bind iron through specialized siderophores (Hermenau *et al*., 2018). Different strains from the species *P. graminis* have shown plant growth-promoting abilities and plant protection against abiotic stresses (Barriuso *et al*., 2008). *P. graminis* is a rhizospheric colonizer of grasses (Viallard *et al*., 1998) and concentrates preferentially at the root surface, but also shows an endophytic lifestyle (Castanheira *et al*., 2016).

To assess the influence of plant growth substrate on the genes important for OAS925 to colonize the plant rhizosphere, we grew *B. distachyon* plants in six different growth substrates. To facilitate this task, we used sterilized substrates, to avoid the effect that other microbes could have on fitness genes. Using RB-TnSeq (Wetmore *et al*., 2015), we showed how genes involved in plant colonization are influenced by plant growth substrates. From the 382 rhizosphere colonization genes that we identified in one or more substrates, only 34 were shared across all 6 conditions, while the remaining 91.1% of the identified fitness genes were impacted by type of substrate. To facilitate follow-up studies, we generated a collection of arrayed transposon mutants in OAS925. Using these individual mutants, we validated our results by assaying their colonization phenotype with plants grown in the different substrates. Additionally, we found that a group of genes that were influenced by plant growth substrate in strain OAS925 were also impacted by substrate type in *P. phytofirmans* PsJN (Sessitsch *et al*., 2005), *Variovorax* sp. OAS795 (Coker *et al*., 2022), and *Rhizobium* sp. OAE497 (Coker *et al*., 2022). Globally, our data shows the importance of taking plant growth substrate into consideration in plant-bacteria interaction studies, especially those that aim to identify fitness genes to design more effective rhizosphere management strategies, engineer strains to improve their plant colonization efficiency, or edit microbial communities *in situ* (Mueller & Sachs, 2015; Rubin *et al*., 2022; Compant *et al*., 2024).

## 2. Material and Methods

### 2.1 Bacterial strains and growth media

Strain OAS925 was originally isolated from a switchgrass (*Panicum virgatum*) field in Oklahoma (United States) (Ceja-Navarro *et al*., 2021; Coker *et al*., 2022). It was routinely cultured in R2A at 30°C with rotary shaking (180 rpm) when needed. The genome of OAS925 was sequenced by the Department of Energy Joint Genome Institute (JGI) using the PacBio Sequel II instrument, and is available at NCBI (accession GCA_040543975.1). Using the Genome Taxonomy Database toolkit v1.1.0 (Chaumeil *et al*., 2020; Parks *et al*., 2022), we determined that OAS925 (NCBI taxonomy ID: 2663827) was most closely related (ANI = 98.5) to the type strain of *P. graminis*. Single OAS925 mutants were grown in R2A with corresponding antibiotics. All *Escherichia coli* strains were routinely cultured in LB at 37°C with 180 rpm shaking when needed. Red fluorescent protein (RFP)-tagged strains were generated introducing plasmid pGingerBK-J23100 (Pearson *et al*., 2023) by conjugation and selecting RFP-transformants by plating on rich media (R2A or LB) supplemented with kanamycin. Unless stated otherwise, antibiotics were used at the following final concentrations: kanamycin (Km) 10 µg/mL, carbenicillin (Cb) 50 µg/mL. When appropriate, the antifungal compound cycloheximide (Cx) was added at 200 µg/mL. If required, diaminopimelic acid (DAP) was added to a final concentration of 300 µM.

### 2.2 Observation of plant colonization by confocal microscopy

To enable microscopic observation of OAS925 bacterial cells colonizing plant roots we used a RFP-tagged strain. *B. distachyon* Bd21-3 seeds were sterilized and germinated (**Supporting Information Note S1**). Seedlings were transferred to Imaging EcoFABs (Jabusch *et al*., 2021) filled with 0.5X Murashige & Skoog (Murashige & Skoog, 1962) (MS) basal salts (MSP01, Caisson Laboratories, United States). 15-day old plants were inoculated with 10^6^ colony forming units (CFUs) of the RFP-tagged OAS925 strain. Seven days after inoculation, plant colonization was observed using a Zeiss LSM 710 confocal microscope system (Zeiss, Germany).

### 2.3 Construction of a barcoded transposon library in *P. graminis* OAS925

The *P. graminis* OAS925 RB-TnSeq library was constructed via conjugation with *E. coli* WM3064 harboring the pHLL250 *mariner* transposon vector library (strain AMD290), which was previously built via Golden Gate assembly using the magic pools approach (Liu *et al*., 2018b); see **Note S2** for methodology details. The final mutant library was named *Paraburkholderia*_OAS925_ML2. To map the genomic locations of the transposon insertions and link these insertions to their associated DNA barcodes, we used a variation of previously described TnSeq protocol (Wetmore *et al*., 2015), where we use two rounds of PCR to selectively enrich for transposon junctions (Rubin *et al*., 2022).

### 2.4 Arrayed collection of OAS925 transposon mutants

To obtain single transposon mutants of *Paraburkholderia_*OAS925_ML2, we assembled an arrayed collection of individual transposon mutants (**Supporting Information Table S1**). To achieve this, we grew one aliquot of the library to OD600 of 0.2 and plated different dilutions on R2A media supplemented with 10 µg/mL Km. Plates were incubated at 30°C for 72 h to let visible colonies develop. 6,912 isolated individual mutants (colonies) were picked into the wells of seventy-two 96-deep well plates with 400 µL R2A with 10 µg/mL Km per well and allowed to grow to saturation at 30°C. For an additional 12 plates (plates 73 to 84), we diluted the library and inoculated the wells such that we added ∼10-100 mutants per well. We did this to generate a larger number of mapped mutants, although mutants in plates 73 to 84 require additional screening to identify the mutant of interest. For all plates, the cultures were mixed with glycerol to a final concentration of 15% v/v for cryopreservation, aliquoted into multiple copies, and stored at -80°C. A well-plate multiplexing strategy where we 1) pooled the cultures from all 96 wells of each plate individually and 2) pooled the same well from all 84 plates combined with barcode sequencing was used to locate the position of each transposon mutant within the arrayed collection (Shiver *et al*., 2021; Arjes *et al*., 2022).

### 2.5 Plant growth conditions

*B. distachyon* Bd21-3 seeds were sterilized and germinated as described before (**Note S1**). *Medicago truncatula* A17 and *Arabidopsis thaliana* Col-0 plants were prepared by surface-sterilization of seeds using standard protocols and germination in 1% w/v water-agar plates in a 130 µmol/m^2^ s^−1^ 16-h light/8-h dark regime at 26°C for 3 days.

Seedlings were transferred to Magenta GA-7 plant culture boxes with vented lids (Caisson Laboratories, United States) filled with different plant growth substrates (**Supporting Information Fig. S1**). Each Magenta box was filled with a volume of ∼100 mL of the different substrates (quartz sand, clay, 0.5X MS liquid, 0.5X MS agar, potting mix and liquid soil extract). For preparation details see **Note S3,** for exact number of replicates per condition see section 2.7. Plants were placed randomly and maintained in a 130 µmol/m^2^ s^−1^ 16-h light/8-h dark regime at 26°C.

### 2.6 Plant growth and rhizosphere metabolite composition across different substrates

In order to determine the metabolite composition of the various plant/substrate conditions, we grew non-inoculated plants prepared as described above and harvested root exudates and rhizosphere metabolites for hydrophilic liquid interaction chromatography and tandem mass spectrometry (LC-MS/MS) analysis. Conditions included a medium-only control (0.5X MS), plants in liquid substrates (0.5X MS liquid and liquid soil extract), and plants in wetted solid substrates (0.5X MS agar, quartz sand, clay, and potting mix). Four weeks after transplanting the seedlings into the substrates, plants were harvested with tweezers and root exudates and rhizosphere metabolites were collected (**Note S4**). Separated roots and shoots were weighed after drying in an oven at 55°C for 48 h followed by cooling to room temperature. Four replicates were used per condition. For all conditions except 0.5X MS liquid, the extractions include metabolites that were available from both root exudation and carbon from plant growth substrates; we will refer hereafter to these "plant/substrate metabolite mixtures" as “rhizosphere metabolites”.

Frozen samples were lyophilized and prepared for LC-MS (**Note S4**). Metabolomics data were processed using Metabolite Atlas software to extract ion chromatograms and peak heights for each metabolite (Yao *et al*., 2015). Metabolites were annotated by comparison of sample m/z, retention time, and MS2 data with an in-house library of authentic reference standards (**Table S2**). Extraction controls and media controls were used to identify and remove contaminants. The raw metabolomics data were deposited to the Global Natural Products Social Molecular Networking (https://gnps.ucsd.edu) data repository (https://gnps2.org/status?task=a4cc328c16d94474af0af7eeacb3155e).

### 2.7 RB-TnSeq fitness assays

Plant and *in vitro* fitness assays were conducted with the *Paraburkholderia*_OAS925_ML2 transposon library. For each batch of experiments, one aliquot of the glycerol stock containing the RB-TnSeq library was thawed and inoculated in 25 mL fresh R2A with 10 µg/mL Km and grown for approximately 13 h at 30°C and 180-rpm shaking until the culture reached OD600 of 1.3-1.6. Time0 (input) samples were collected from the recovered library by centrifuging 1 mL aliquots and freezing at -20°C until DNA purification. The remaining cells were then washed twice with a chemically-defined media without a carbon source, RCH2_defined_noCarbon (Price *et al*., 2018), unless noted otherwise. A summary of all genome-wide fitness assays performed in this study with OAS925, which include 8 plant conditions (6 with *Brachypodium*, 1 with *Medicago*, 1 with *Arabidopsis*), 2 outgrowth controls and 110 *in vitro* conditions, as well as the number of replicates used per condition, can be found in **Fig. 1** and **Table S3**.

**Figure 1.**
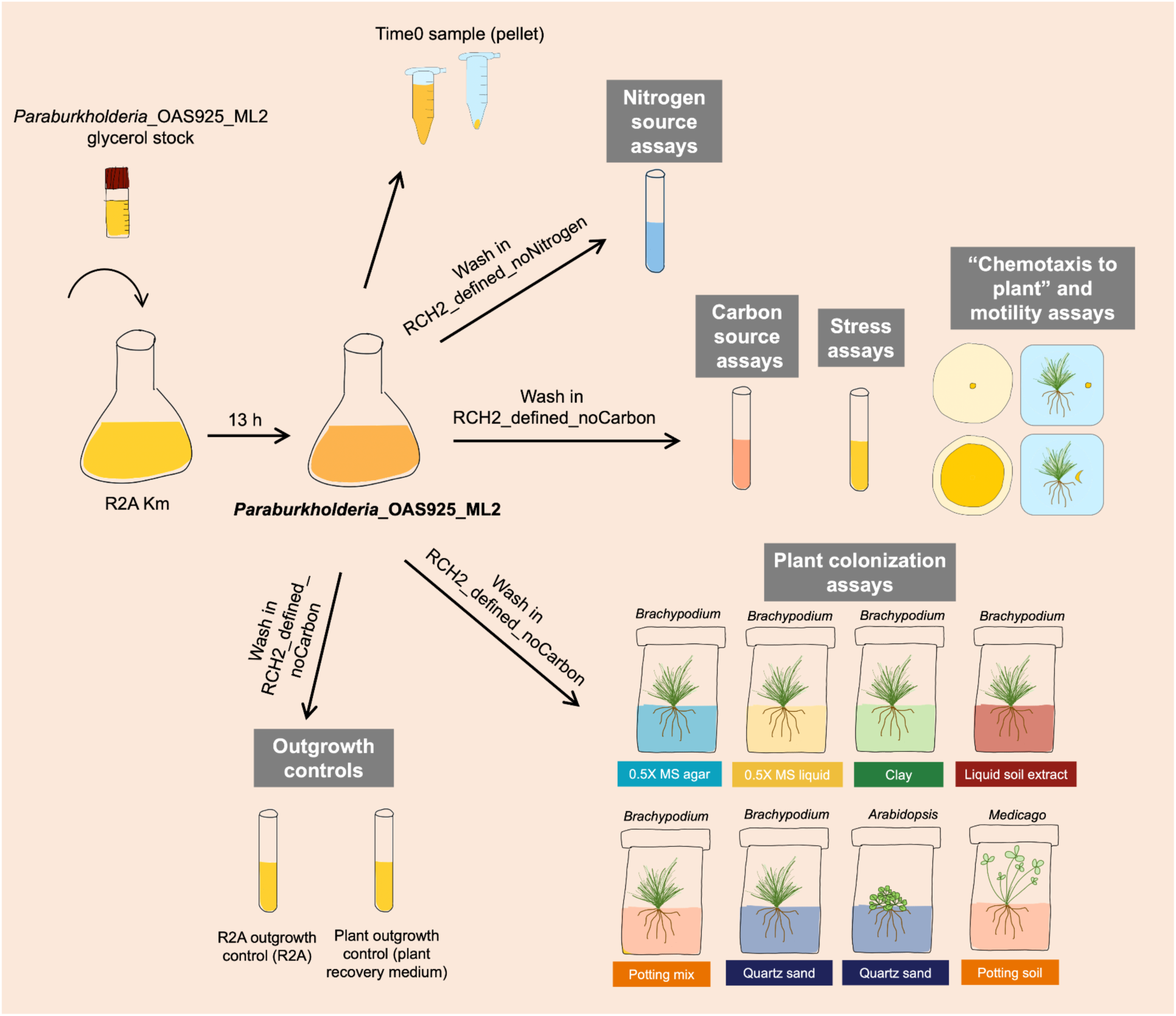
Overview of the RB-TnSeq *in vitro* and *in planta* fitness assays conducted with *Paraburkholderia*_OAS925_ML2. Time0 (input) samples are the common control for all experimental conditions. For the plant colonization assays, the washed library was inoculated in each 15-day old plant by injecting directly on the substrate with a sterile pipette in 6-7 different spots at ∼1 cm around the plant stem. A summary of the conditions tested can be found in Table S3.

### **a)** Plant colonization fitness assays

The *Paraburkholderia*_OAS925_ML2 library was washed twice with RCH2_defined_noCarbon. Cells were then harvested by centrifugation (3,000 *g* for 3 min), and resuspended in RCH2_defined_noCarbon to reach OD600 = 1. Then, 40 µL (∼10^7^ CFU) were diluted in 5 mL of 0.5X MS and the whole volume was inoculated in each 15-day old plant by injecting directly on the substrate with a sterile pipette in 6-7 different spots at ∼1 cm around the plant stem. The substrate was stirred with a sterile spatula around the plant to homogenize as much as possible the bacterial inoculum.

Magenta culture boxes were placed randomly and maintained in a 130 µmol/m^2^ s^−1^ 16-h light/8-h dark regime at 26°C for three weeks. Every 7 days, substrates (except 0.5X MS liquid, 0.5X MS agar and liquid soil extract) were watered with 5 mL of sterile water and stirred with a sterile spatula. In total, between 4 and 9 replicates were conducted for each *Brachypodium* condition, and 2 replicates (each a pool of three plants) for *Medicago* and *Arabidopsis* (**Table S3**). Replicates were conducted on separate days.

Twenty-one days after the inoculation with *Paraburkholderia*_OAS925_ML2, samples were harvested as follows. Plants were gently uprooted, and the aerial part (root/shoot junction) was cut using sterile scissors and disposed of. Isolated roots were collected in 50-mL falcons containing 10 mL of plant recovery medium (R2A supplemented with 10 µg/mL Km, 200 µg/mL Cx, and 0.01% v/v Tween 20) and vortexed at high speed for 2 minutes. This sample was referred to as the “rhizosphere”. After vortexing, roots were discarded and the liquid content was transferred to glass tubes. Samples were then incubated for 35 h at 30°C and 180-rpm shaking. Two-mL samples from all cultures were harvested. After centrifugation (3,000 *g* for 3 min), pellets were stored at -20°C until DNA purification.

As outgrowth controls, 40 µL of the washed library were inoculated on both R2A (R2A outgrowth control) and plant recovery medium (plant outgrowth control), respectively) and grown for 35 h at 30°C with 180-rpm agitation. Two-mL samples were collected, centrifuged and frozen at -20°C until further DNA purification. Ten replicates were performed for each control (**Table S3**).

### b) *In vitro* fitness assays

*In vitro* fitness assays (**Table S3**) were conducted in defined media or rich media, depending on the type of experiment. For all assays except the nitrogen source assays, the mutant library was washed using RCH2_defined_noCarbon (**Fig. 1**). For nitrogen source assays, cultures were washed using RCH2 media without NH4Cl and supplemented with 20 mM sodium DL-lactate (referred to as RCH2_defined_noNitrogen).

Growth assays (e.g. carbon metabolism, nitrogen metabolism, and stressors/inhibitors) were performed as previously described (Wetmore *et al*., 2015; Price *et al*., 2018) either in glass tubes (10-mL per tube) or 96 deep-well microplates (0.8 mL per well) (see **Note S5** for details). Biofilm assays were conducted as previously described (O’Toole, 2010).

For swimming motility assays, the mutant library was inoculated into the center of 0.25% w/v agar R2A plates, and “outer” samples with motile cells were removed with a sterile razor after 24 h. “Inner” samples of cells from near the point of inoculation were also removed (Price *et al*., 2018). For “chemotaxis to plant root” assays, the mutant library was inoculated at ∼1 cm from 21-old-day plants that were transferred the day before inoculation onto 0.5X MS plates with 0.25% w/v agar (**Fig. S2**). After 24 h, outer samples with motile cells were removed with a razor. Overnight outgrowths of inner and/or outer samples were conducted in 10 mL of R2A, centrifuged, and pellets were kept at -20°C until DNA purification.

### 2.8 DNA isolation, library preparation, sequencing, and fitness value calculation

DNA from frozen pellets was isolated using the DNeasy Blood & Tissue Kit (Qiagen, Germany) according to manufacturer’s instructions. Purified RNA-free DNA was used as a template for DNA barcode PCR amplification using previously described PCR conditions (Wetmore *et al*., 2015) (**Note S6**). Strain and gene fitness calculations were done using the computation pipeline developed by Wetmore et al. (Wetmore *et al*., 2015); code available at bitbucket.org/berkeleylab/feba/src/master. Fitness values for each gene were calculated as the log2 of the ratio of relative barcode abundance for mutants in that gene following library growth in a given condition (i.e. plant colonization) divided by relative abundance in the Time0 (input) sample. A t-like statistic was also determined for every gene in each experiment, which takes into account the consistency of the fitness of all the mutants of that gene to assess if the fitness value is reliably different from 0 (Wetmore *et al*., 2015). To avoid considering weak colonization fitness values, which may not be biologically meaningful, we focused on genes that had |fitness| ≥ 1, with an associated |t| ≥ 3. The majority of the fitness data from this study is publicly available at the Fitness Browser website (https://fit.genomics.lbl.gov) (Price *et al*., 2018) for comparative analysis. Some experiments did not pass our thresholds for inclusion in the Fitness Browser; the data from these experiments is available in the corresponding tables (see further below, **Table S7, Table S14, Table S15**, and **Table S16**).

### 2.9 Fitness data analysis

Analysis of gene fitness values was done in R (R Core Team, 2017). Fitness values and t scores were combined across replicates by averaging them; with the expectation that doing this for t values is conservative. For a gene to be considered as involved in fitness in a certain condition, both average fitness value (|fitness| ≥ 1) and average t value (|t| ≥ 3) criteria needed to be met. Genes that showed a fitness effect in the R2A outgrowth control and/or plant outgrowth control were removed from the analysis of plant colonization genes. Graphs and heatmaps were plotted in R using the *ggplot2* package (Wickham, 2016). Principal component analysis (PCA) plots of experiments were generated from the matrix of gene fitness values using the function *prcomp*.

### 2.10 Confirmation of gene functionality by testing individual mutants in plants

We used individual mutants from the arrayed collection to verify our genome-wide RB-TnSeq data. All cultures were started from individual colonies, and the correct barcoded mutant was confirmed by PCR and Sanger sequencing. To confirm gene function and validate the RB-TnSeq results, we performed colonization assays with individual transposon mutants that exhibited different fitness in at least two growth substrates, for which we used plants grown in those substrates. *Brachypodium* plants were prepared as described above. Briefly, overnight bacterial cultures were washed twice with RCH2_defined_noCarbon and adjusted to OD600 = 1. Then, each plant was inoculated with a mixture of 40 µL of the wild-type and 40 µL of the mutant strain, both resuspended in 5 mL of 0.5X MS and inoculated in each 15-day old plant by injecting directly on the substrate in 6-7 different spots at ∼1 cm around the plant stem. Then, the substrate was stirred with a sterile spatula around the plant to homogenize as much as possible the bacterial inoculum. Plants were incubated for three weeks as explained above. Then, rhizosphere samples were harvested as detailed before, and CFU of wild-type and mutant populations were determined in selective media by comparing colony counts on R2A versus R2A with Km plates. CFU of each mutant was determined on R2A Km, while CFU of wild-type strain was obtained after subtracting the number of colonies in R2A Km from the number of colonies in R2A. Competitive index (CI) values were calculated using at least 3 replicates per condition following the formula (CFU mutant/CFU wild-type) described by Macho et al. (Macho *et al*., 2010); CI < 1 means the mutant is less fit than the wild-type, CI > 1 means the mutant is more fit than the wild- type. As controls, we also performed competition assays in R2A media with the wild-type strain in the same 1:1 proportion.

### 2.11 Evaluation of the impact of plant growth substrate on the colonization fitness of other plant-associated bacteria

To evaluate the influence of plant growth substrate on the colonization fitness of other plant-related bacteria, we used RB-TnSeq libraries of *P. phytofirmans* PsJN, *Variovorax* sp. OAS795 (NCBI taxonomy ID: 3034231; NCBI RefSeq assembly GCF_040546685.1; most closely related to *V. paradoxus*) and *Rhizobium* sp. OAE497 (NCBI taxonomy ID: 2663796; NCBI RefSeq assembly GCF_040546565.1; most closely related to *R. endophyticum*). PsJN is a well-established model for plant-associated endophytic bacteria that is able to colonize a wide range of plants (Sessitsch *et al*., 2005). OAS795 is a strain originally isolated from the same field as OAS925 (Coker *et al*., 2022) which is able to quickly dominate the population of unplanted and *Brachypodium*-planted soil inoculated with a 17-member SynCom, but is not as competitive in *B. distachyon* root and rhizosphere (Lin *et al*., 2023). OAE497 was isolated in Oklahoma (Coker *et al*., 2022) from the same field as OAS795 and OAS925, and can dominate the population of *Brachypodium* rhizosphere inoculated with a 17-member SynCom, but to a lesser extent than OAS925 (Lin *et al*., 2023). We observed plant colonization of PsJN, OAS795, and OAE497 by confocal microscopy as explained in sections 2.1 and 2.2.

The RB-TnSeq mutant library of PsJN, named BFirm_ML3, was previously described (Price *et al*., 2018). The OAS795 and OAE497 RB-TnSeq mutant libraries were constructed for this study (**Note S7**). Plant colonization experiments were conducted with BFirm_ML3, *Variovorax*_OAS795_ML2, and *Rhizobium*_OAE497_ML4 as follows. *Brachypodium* plants were grown in quartz sand and 0.5X MS agar as described in section 2.5; up to 5 replicates were used per condition. *In vitro* assays were also conducted with *Variovorax*_OAS795_ML2 with 50 different carbon sources. In total, 117 fitness assays (see summary in **Table S4**) were conducted with BFirm_ML3, *Variovorax*_OAS795_ML2 and *Rhizobium*_OAE497_ML4. Fitness assays, DNA isolation, library preparation, sequencing, fitness value calculation, and data analysis were done as explained for OAS925. In order to allow comparison of the fitness data of PsJN, OAS795, and OAE497 with the OAS925 fitness data we used orthologue data inferred with OrthoFinder (Emms & Kelly, 2019).

## 3. Results

### 3.1 *P. graminis* OAS925 efficiently colonizes *B. distachyon* roots

*P. graminis* OAS925 was isolated from the rhizosphere of switchgrass grown on marginal soil (Ceja-Navarro *et al*., 2021; Coker *et al*., 2022). To determine if this strain is capable of colonizing *Brachypodium* roots, we generated an RFP-tagged strain and observed colonization using Imaging EcoFABs, fabricated ecosystems for non-destructive high-resolution *in situ* examination of microbial establishment on plants (Jabusch *et al*., 2021). As observed by confocal microscopy at 20X and 40X amplification, OAS925 colonizes very efficiently the rhizosphere and roots of *B. distachyon* plants seven days after inoculation (**Fig. 2**).

**Figure 2.**
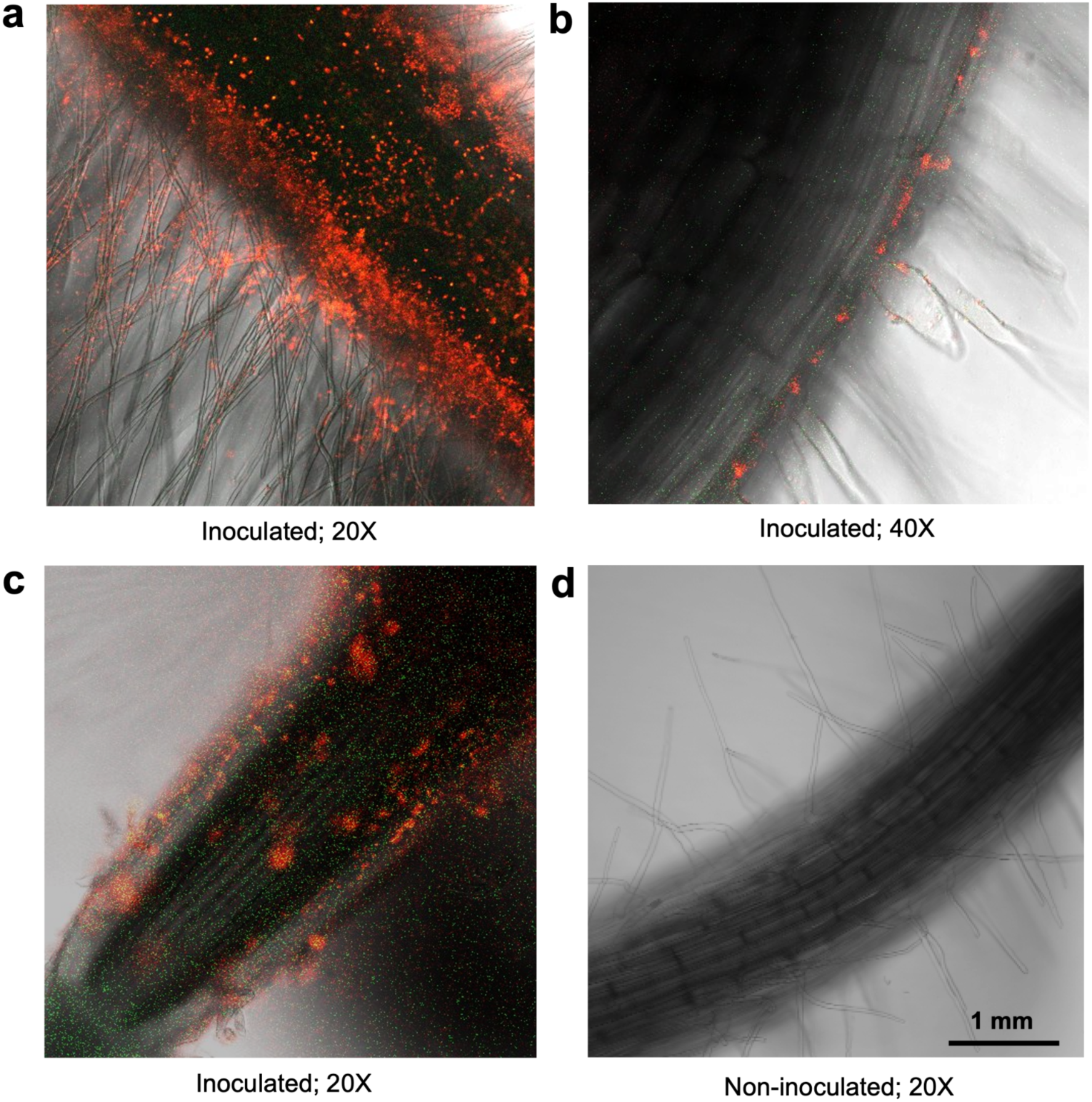
Confocal microscopy of *Brachypodium distachyon* Bd21-3 roots colonized by a RFP-tagged strain of *Paraburkholderia graminis* OAS925. Colonization was observed 7 days after inoculation in plants grown in Imaging EcoFABs at 20X (a, c, d) and 40X (b) amplification using a Zeiss LSM 710 confocal microscope system. Overlay of fluorescence and bright-field microscopy is shown. a, b, c) Inoculated plants. d) Non- inoculated plants.

### 3.2 Type of substrate impacts *B. distachyon* growth parameters and rhizosphere metabolite composition

To assess the impact of the different growth substrates (**Fig. S1a**) on plant growth parameters, we grew *B. distachyon* in sterile substrates without the addition of strain OAS925, and we weighed root and shoot biomass 36 days after germination. The results show that root biomass was very low in potting mix compared to the other 5 substrates, and the root biomass in liquid soil extract, quartz sand, and clay was slightly higher than in the 0.5X MS agar and 0.5X MS liquid experiments (**Fig. 3a**). Although we did not systematically evaluate this, root morphology, and more specifically the amount of root hairs, was observed to be different across some of the substrates used, in particular between 0.5X MS liquid and liquid soil extract (**Fig. S1b**). Shoot weight was the lowest in potting mix as well, while reaching the highest values in 0.5X MS agar, 0.5X MS liquid, and liquid soil extract (**Fig. 3b**).

**Figure 3.**
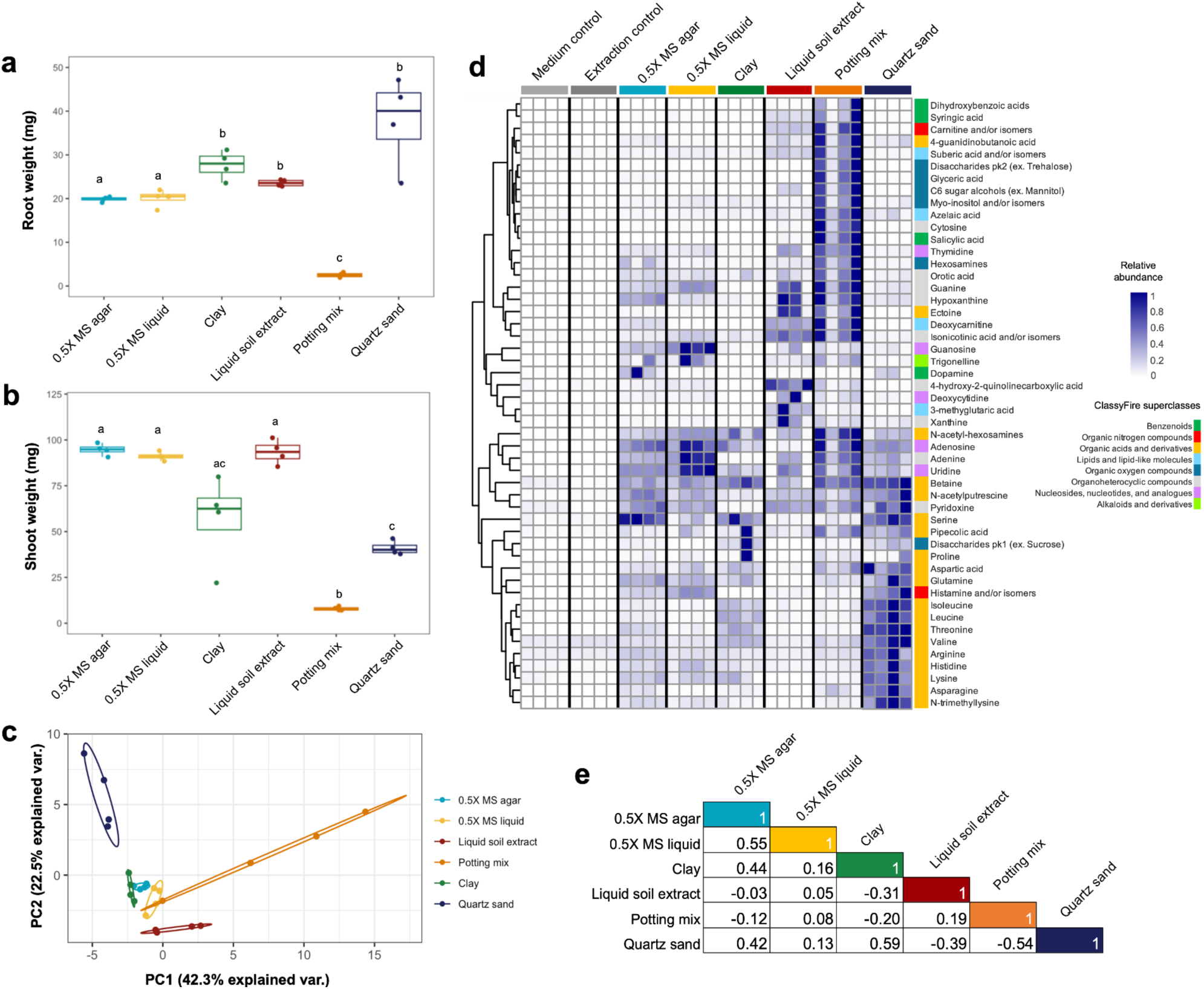
Impact of plant growth substrate on *Brachypodium distachyon* Bd21-3 growth and root exudate composition. **a**) Root and **b**) shoot dry weight of plants grown in diverse substrate types. Different letters above the bars indicate that the means differ significantly (*P* &lt; 0.05) across conditions. **c**) Principal component analysis (PCA) of the rhizosphere metabolites present in plants grown in different types of growth substrate; relative abundance of peak height is used. **d**) Clustered heatmap of rhizosphere metabolite composition across plants grown in different substrates, as well as two controls; relative abundance of peak height is shown. Superclass according to ClassyFire chemical classification is shown (Djoumbou Feunang *et al*., 2016). **e**) Pearson pairwise correlation of average rhizosphere metabolite composition across different plant growth substrates.

To characterize the rhizosphere metabolites which were present in the different conditions, we performed metabolomics analysis on the rhizosphere environment of non-inoculated plants grown in the different types of substrates. Detected metabolites were consistent with metabolites identified in our previous studies (Sasse *et al*., 2019) and were found to vary in abundance across conditions (**Fig. 3c**, **Fig. 3d**, **Fig. S3**, **Table S5**). Notably, we observed that some compound classes (ClassyFire superclasses) (Djoumbou Feunang *et al*., 2016) were more abundant in certain conditions. We for example observe that benzenoids (e.g. salicylic acid, syringic acid, dihydroxybenzoic acids) are more abundant in plants grown in potting mix, while organic acids and derivatives (e.g. asparagine, histidine, lysine, valine) were in higher abundance in the quartz sand condition (**Fig. 3d**, **Fig. S3**, **Table S5**). As observed by Pearson pairwise correlation of average rhizosphere metabolite composition, quartz sand and clay, followed by 0.5X MS agar and 0.5X MS liquid, had the most correlated metabolite profiles, while the lowest correlation value was between quartz sand and potting mix (**Fig. 3e**).

### 3.3 Bacterial plant rhizosphere colonization genes are influenced by plant growth substrate

With the purpose of identifying plant colonization fitness genes in strain OAS925, we generated an RB-TnSeq *mariner* mutant library that we named *Paraburkholderia*_OAS925_ML2. Mutagenesis and subsequent TnSeq resulted in a pool of 402,743 mutant strains with mapped insertions and unique barcodes at 128,471 different locations distributed across the genome (**Fig. 4a**) at an average rate of 1 insertion location every 56 bp. Of the 6,579 predicted genes in the genome of OAS925, we generated mutations in 5,667 genes (86.1% of the genome). The remaining 912 genes do not have gene fitness data because they have few or no mutants in the library (many are likely essential genes), or mutants in these genes had a low number of reads in the Time0 samples (**Table S6**).

**Figure 4.**
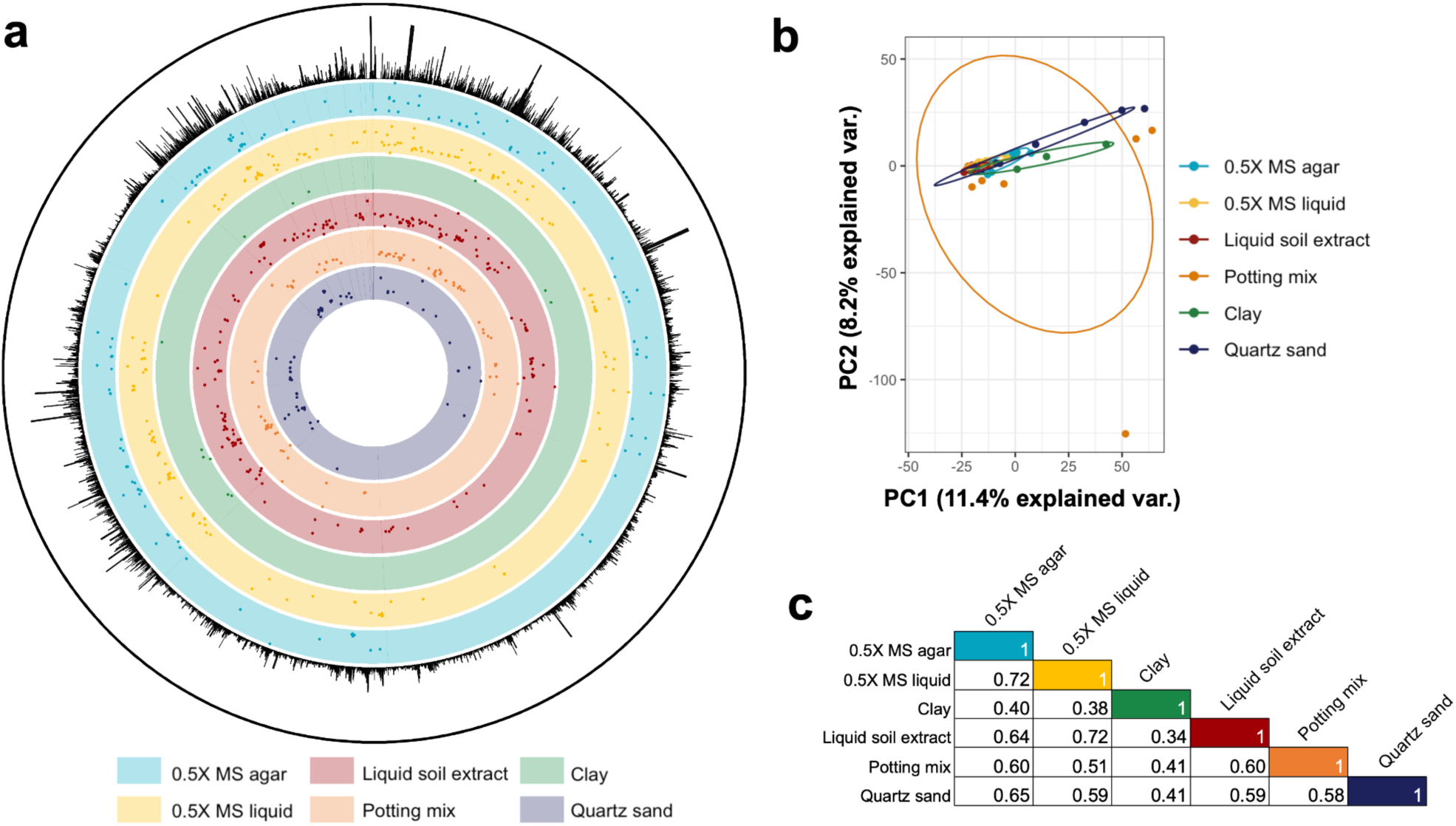
Influence of plant growth substrate type on bacterial fitness for plant colonization. **a**) Genome-wide map of *Paraburkholderia*_OAS925_ML2 plant rhizosphere colonization genes. Outer to inner tracks: total sequencing reads per gene from TnSeq mapping data; rhizosphere colonization genes (|fitness| ≥ 1 and |t| ≥ 3) across plant growth substrates (light blue, 0.5X MS agar; yellow, 0.5X MS liquid; green, clay; red, liquid soil extract; orange, potting mix; dark blue, quartz sand). **b**) Principal component analysis of *Paraburkholderia*_OAS925_ML2 rhizosphere colonization genes in different substrate types. Plot generated using gene fitness values for 5,667 genes. **c**) Pearson pairwise correlation of average plant fitness data across different growth substrates.

To evaluate the influence of plant growth substrate on rhizosphere colonization fitness genes in OAS925_ML2 we designed *B. distachyon* rhizosphere colonization assays using 6 different types of substrates. The library was also tested in the plant species *A. thaliana* and *M. truncatula* and in 110 *in vitro* conditions in order to facilitate further characterization of rhizosphere colonization genes. We used sequenced barcode read counts to quantify the representation of mutants in each experiment and compared barcode frequencies across conditions. In total, we performed 298 genome-wide fitness experiments (**Table S3**). Fitness values for each gene were calculated and averaged across each condition as the log2 of the ratio of relative barcode abundance for mutants in that gene following library growth in a given condition divided by relative abundance in the Time0 sample (see section 2.8) (**Table S7**, **Table S8**). A negative fitness means that mutants in the gene are depleted in that given condition; these are called “important” genes, that when mutated, result in reduced colonization ability. A positive fitness means that mutants in the gene are enriched in that condition; these are called “detrimental” genes, that when mutated, result in improved colonization ability.

Genes significantly contributing to colonization fitness were distributed throughout the OAS925 genome (**Fig. 4a**), with many clustering together. Since 99 genes (**Table S9**) had a mild fitness defect or advantage (|fitness| ≥ 0.75 and |t| ≥ 4) in the outgrowth controls (i.e. R2A and plant recovery medium), they were not considered in our plant fitness analysis. In total, 382 genes (5.8% of all OAS925 genes; **Table S10**) had a significant phenotype (|fitness| ≥ 1 and |t| ≥ 3) in *B. distachyon* rhizosphere colonization in at least one plant growth substrate type. To explore the physiological functions of these 382 rhizosphere colonization genes in more detail, we compared our data to RB-TnSeq results of the same insertion mutant library tested under a large number distinct plant-related and *in vitro* conditions. The latter included colonization of other plant species (*A. thaliana* and *M. truncatula*), metabolism of 76 carbon and 6 nitrogen sources commonly found in root exudates, tolerance to 20 different stress conditions frequently found in the plant environment (including pH, oxygen reactive species, plant hormones, and coumarins), motility, and chemotaxis to plant roots. Although the complexity of individual phenotypes measured by these *in vitro* experiments is considerably lower than that of rhizosphere colonization processes, these assays can be used to rapidly assess many metabolic or stress responsive functions. Despite our additional screening efforts, 54 genes (**Table 1, Table S11**) from the 382 rhizosphere colonization genes were uniquely involved in plant colonization (|fitness| ≥ 1 and |t| ≥ 3 in at least one growth substrate) and did not significantly (|t| ≤ 3) contribute to fitness in any of the *in vitro* conditions tested (**Table S11**).

**Table 1.**
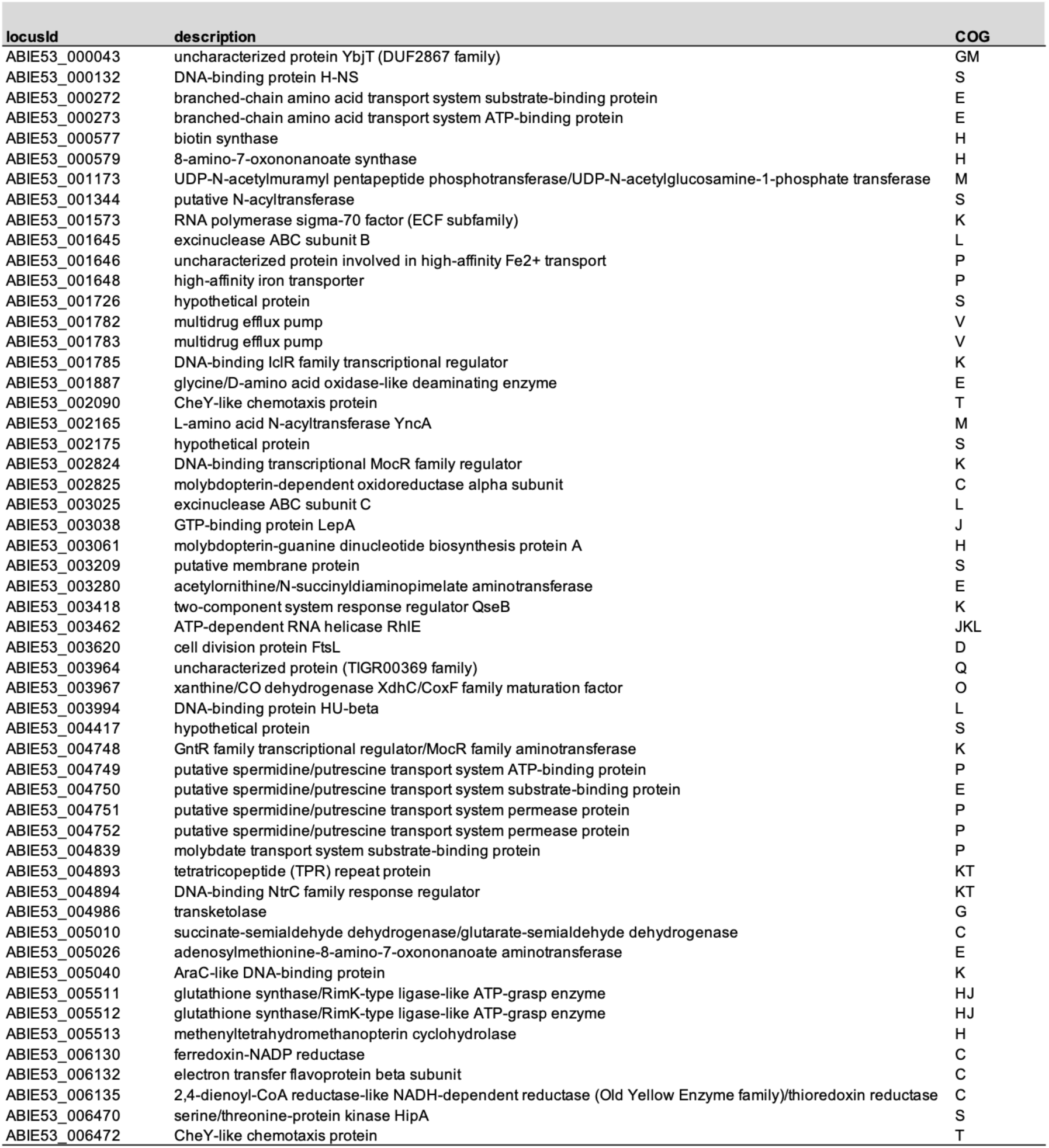
List of *Paraburkholderia*_OASA925_ML2 genes involved uniquely in *Brachypodium distachyon* Bd21-3 rhizosphere colonization.

***OAS925 genes shared across substrates.*** Whereas the majority of the 382 rhizosphere colonization genes that we identified in OAS925 only had a significant phenotype (|fitness| ≥ 1 and |t| ≥ 3; **Table S10**) in one or a few of the substrates tested, a group of genes was needed across all growth substrates. Using the criteria |fitness| ≥ 1 and |t| ≥ 3 in all plant growth substrates, 6 genes were identified as shared across all substrates. We also reasoned that, if a gene had |fitness| ≥ 1 and |t| ≥ 3 in at least one plant growth substrate, and |fitness| ≥ 1 and |t| = 1-3 in the other types of substrates, then that gene might be required for colonization across all plant growth substrates (despite the relaxed t value); and therefore we focused on the 34 genes (8.9% of the total identified rhizosphere genes) meeting that criteria (**Table 2, Table S12**). We examined the predicted functions of the identified 34 “core” colonization genes based on clusters of orthologous groups (COG classification) of protein annotations, which we obtained with the eggNOG-mapper v2 tool (Cantalapiedra *et al*., 2021), and found that 7 of these genes are related to amino acid transport and metabolism (COG class E), six to transcription (COG class K), and six to energy production and conversion (COG class C).

**Table 2.** List of *Paraburkholderia*_OASA925_ML2 genes involved in *Brachypodium distachyon* Bd21-3 rhizosphere colonization across all types of substrates.

| locusId | description | COG |
| --- | --- | --- |
| ABIE53_000046 | putative MarR family transcription regulator | K |
| ABIE53_000413 | methionine biosynthesis protein MetW | Q |
| ABIE53_000414 | homoserine O-acetyltransferase | E |
| ABIE53_000423 | DnaK suppressor protein | T |
| ABIE53_000650 | thiol-disulfide isomerase/thioredoxin | CO |
| ABIE53_000674 | cytochrome c oxidase subunit 1 | C |
| ABIE53_000678 | cytochrome c oxidase subunit 3 | C |
| ABIE53_000682 | cytochrome c oxidase assembly protein subunit 15 | O |
| ABIE53_000683 | protoheme IX farnesyltransferase | H |
| ABIE53_000759 | 7-cyano-7-deazaguanine reductase | F |
| ABIE53_001228 | phosphoadenosine phosphosulfate reductase | C |
| ABIE53_001947 | CRP-like cAMP-binding protein | K |
| ABIE53_002004 | anti-sigma factor RsiW | K |
| ABIE53_002424 | anti-sigma factor RsiW | K |
| ABIE53_003114 | DNA-binding NtrC family response regulator | K |
| ABIE53_003148 | UDPGlucose 6-dehydrogenase | C |
| ABIE53_003157 | chorismate mutase/prephenate dehydratase | E |
| ABIE53_003158 | phosphoserine aminotransferase | E |
| ABIE53_003306 | ATP-dependent Clp protease ATP-binding subunit ClpA | O |
| ABIE53_003460 | mono/diheme cytochrome c family protein | C |
| ABIE53_003556 | 2-dehydro-3-deoxyphosphogalactonate aldolase | G |
| ABIE53_003638 | ribulose-phosphate 3-epimerase | G |
| ABIE53_003733 | glutamate synthase domain-containing protein 3 | E |
| ABIE53_003734 | glutamate synthase (NADPH/NADH) large chain | E |
| ABIE53_003735 | hypothetical protein | S |
| ABIE53_004801 | 3-isopropylmalate/(R)-2-methylmalate dehydratase large subunit | H |
| ABIE53_004803 | 3-isopropylmalate/(R)-2-methylmalate dehydratase small subunit | E |
| ABIE53_004804 | 3-isopropylmalate dehydrogenase | CE |
| ABIE53_004817 | O-succinylhomoserine sulfhydrylase | E |
| ABIE53_004820 | K<sup>+</sup>-sensing histidine kinase KdpD | T |
| ABIE53_004858 | farnesyl diphosphate synthase | H |
| ABIE53_004866 | RNA polymerase primary sigma factor | K |
| ABIE53_005493 | aspartokinase-like uncharacterized kinase | S |
| ABIE53_006135 | 2,4-dienoyl-CoA reductase-like NADH-dependent reductase (Old Yellow Enzyme family)/thioredoxin reductase | C |

Of the 34 core rhizosphere genes, 31 are important (fitness ≤ 1) for plant colonization, and 3 are detrimental (fitness ≥ 1). Twenty and 29 out of these 34 genes are also involved (|fitness| ≥ 1) in rhizosphere colonization of *A. thaliana* and *M. truncatula* plants, respectively (**Table S12**). To our knowledge, some of these 34 core colonization genes have not been implicated in plant colonization before: ABIE53_005493 (aspartokinase-like uncharacterized kinase), ABIE53_002004 and ABIE53_002424 (anti- sigma factors RsiW), and ABIE53_000046 (putative MarR family transcription regulator). Several other of these 34 core colonization genes have already been described in plant-associated bacteria as involved in colonization; for instance, ABIE53_003556 (galactonate metabolism) (Cole *et al*., 2017), ABIE53_000413-ABIE53_000414 and ABIE53_004817 (methionine synthesis) (Sivakumar *et al*., 2019; Su *et al*., 2021), ABIE53_004801 (leucine biosynthesis) (Torres *et al*., 2022), ABIE53_000678 and ABIE53_000682-ABIE53_000683 (synthesis of respiratory hemes and cytochrome oxidases) (Torres *et al*., 2022), ABIE53_001228 (cysteine biosynthesis) (Georgoulis *et al*., 2021), and ABIE53_000423 (DnaK suppressor protein) (Zhang *et al*., 2019). In total, the 34 core genes only represent 13.8% of the whole number of genes identified in substrates such as liquid soil extract (245 genes), reflecting the importance of considering several plant growth substrates when assessing bacterial plant colonization fitness.

As assessed by the large number of *in vitro* fitness assays conducted with *Paraburkholderia*_OAS925_ML2, most of the 31 important core colonization genes were also involved (|fitness| ≥ 1, |t| ≥ 3) in the metabolism of C (e.g. fumaric acid, cellobiose, shikimic acid, azelaic acid, alanine, lactate, succinate, and pyruvate) and N (e.g. 1- aminocyclopropane-1-carboxylate, histamine, and xanthine) sources; whereas a selection of them were additionally involved in tolerance to different stresses (e.g. p-coumaric acid), motility and chemotaxis to plant roots (**Fig. S4**, **Table S12**). A few genes only had a phenotype in a reduced number of *in vitro* assays. For instance, apart from rhizosphere colonization, ABIE53_005493 (aspartokinase-like uncharacterized kinase) was only important for the metabolism of glycyl-L-glutamic acid (Gly-Glu). Regarding the 3 detrimental core colonization genes, they were also detrimental for the metabolism of some C sources (e.g. D-mannose, L-alanine, L-proline, L-fucose, L-glutamic acid) (**Fig. S4**, **Table S12**). These results mirror the complex nature of the root colonization process.

### OAS925 genes influenced by substrate type

As shown by the results (**Fig. 4**, **Fig. S5**), the fitness data in OAS925 for colonization of *B. distachyon* plants is distinct depending on the growth substrate in which plants were grown. In a principal component analysis (PCA) we observe that the experiments in 0.5X MS agar, 0.5X MS liquid, and liquid soil extract are more reproducible across replicates, and that those in clay, quartz sand, and potting mix are more variable (**Fig. 4b**). The high variability in potting mix may be explained by how poorly the plants grew in this condition (**Fig. 3a**, **Fig. 3b**). As observed by Pearson pairwise correlation of average plant colonization fitness data, 0.5X MS agar and 0.5X MS liquid were the most correlated, while the lowest correlation value was observed between the clay and liquid soil extract conditions (**Fig. 4c**).

In total, 91.1% (348 from 382) of the rhizosphere fitness genes identified in OAS925 are influenced by plant growth substrate (**Table S10**). The total number of genes contributing to plant colonization fitness was different depending on the substrate type, with liquid soil extract having the most identified genes, and clay the least (**Fig. S5a**).

A fraction of the genes was found to be uniquely involved for rhizosphere colonization of plants grown in one substrate. For instance, 78 genes were found to be important (fitness ≤ -1, t ≤ -3) for plants grown in liquid soil extract, but not in the other plant growth substrates (**Fig. S5b**). Interestingly, a few genes showed opposite fitness values in different types of substrates. For example, part of the cluster ABIE53_001141-ABIE53_001163 (coding for glycosyltransferases, lipopolysaccharide transport systems, and hypothetical proteins) has negative average fitness for colonization of plants grown in 0.5X MS agar, quartz sand, and potting mix, but positive average fitness for plants grown in 0.5X MS liquid and liquid soil extract. Genes in this cluster are important for motility, metabolism of amino acids, and acid pH tolerance; and are detrimental for biofilm formation and sugar metabolism (**Table S10**). Another example is ABIE53_003964 (coding for an uncharacterized protein with a predicted thioesterase domain (Mistry *et al*., 2021; Paysan-Lafosse *et al*., 2023)), that had average fitness = -1.30 in clay, whereas in potting mix it had average fitness = 2.76. Other examples include ABIE53_000102 (flagellar basal-body rod protein FlgB), ABIE53_000111 (flagellar biosynthesis GTPase FlhF) (**Fig. 5**), ABIE53_002702 (hypothetical protein), and ABIE53_005512 (glutathione synthase/RimK-type ligase-like ATP-grasp enzyme).

**Figure 5.**
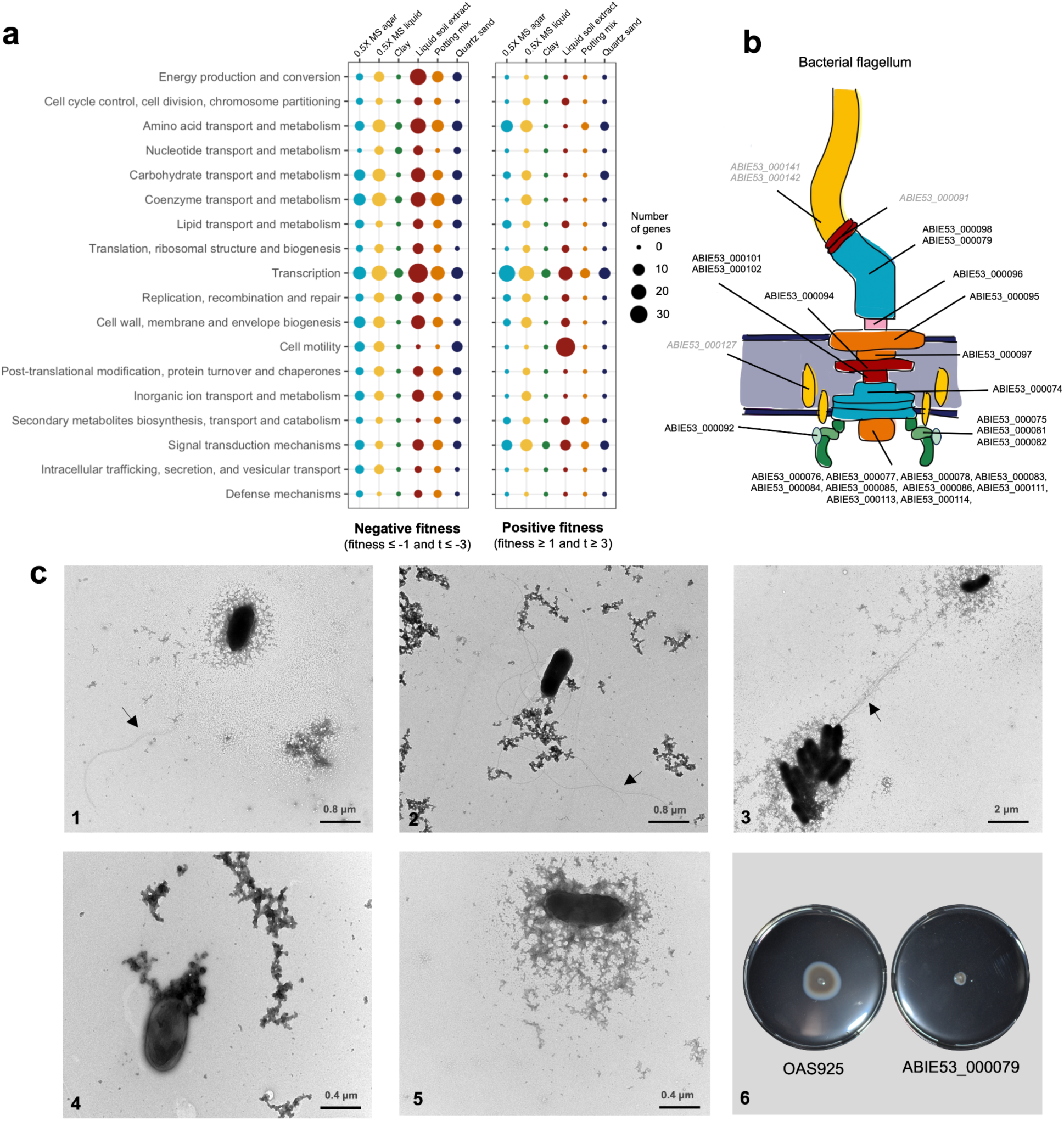
Number of *Paraburkholderia graminis* OAS925 genes per COG category involved in rhizosphere colonization in each of the different plant growth substrates. **a**) Number of genes with a negative fitness or a positive fitness. **b**) Schematic representation of the flagellar structure in strain OAS925. In black font, genes with a positive fitness value in liquid soil extract; in gray and italics, other flagellar genes. **c**) Observation of flagella in wild-type OAS925 (1, 2, 3) and mutant ABIE53_000079 (flagellar hook-length control protein FliK) (4, 5) by transmission electron microscopy (Zeiss LEO906E), after negatively staining overnight bacterial cultures with 1% w/v uranyl acetate. Swimming motility in R2A plates with 0.25% w/v agar 24 h after inoculation.

Not surprisingly, many of the identified genes in a given substrate are clustered together in the OAS925 genome (**Table S10**). For instance, ABIE53_000270- ABIE53_000274, annotated as branched-chain amino acid transport genes, are detrimental for rhizosphere colonization of plants grown in 0.5X MS liquid, 0.5X MS agar, and quartz sand. ABIE53_000295-ABIE53_000307 (related to the general secretion pathway) are important for colonization in 0.5X MS liquid and 0.5X MS agar. Another example is the cluster ABIE53_003398-ABIE53_003400, encoding a predicted zinc/manganese ABC transporter, that only has a negative fitness in plants grown in potting mix and liquid soil extract. Other clusters of genes appear important only in one or a few plant growth substrates, such as ABIE53_004748-ABIE53_004752, involved in spermidine/putrescine transport, and ABIE53_005511-ABIE53_005515, related to the tetrahydromethanopterin and methanofuran pathway, that are identified as detrimental only for colonization of plants grown in 0.5X MS liquid. Other examples are ABIE53_000577-ABIE53_000579 (biotin synthesis), ABIE53_003440 (cyclic pyranopterin phosphate synthase, related to molybdenum cofactor biosynthesis), ABIE53_001075 (hypothetical protein with homology to extracellular matrix protein PelA), ABIE53_002104 (phytoene synthesis), ABIE53_002867 (2-isopropylmalate synthase), ABIE53_004324 (LuxR family quorum-sensing transcriptional regulator), and ABIE53_002805 (two-component system nitrogen regulation response regulator GlnL). These results, along with the observation that many of the genes identified in our screen are involved in processes well known to be vital to colonization of plants (e.g. motility, carbohydrate utilization, amino acid synthesis) are consistent with the notion that fitness values from our genome-wide colonization study reflect valid, biologically relevant genes and pathways.

To examine if different cellular processes are more important for plant colonization in particular substrates, we examined the COG classifications of the identified genes. We found that processes such as energy production and conversion (COG category C), amino acid transport and metabolism (category E), carbohydrate transport and metabolism (category G), coenzyme transport and metabolism (category H), transcription (category K) and cell wall, membrane and envelope biogenesis (category M), among others, appear to be more important (fitness ≤ -1, t ≤ -3) for rhizosphere colonization of plants grown in any substrate compared to clay (**Fig. 5a**). Genes related to motility (category N) were detrimental (fitness ≥ 1, t ≥ 3) only for colonization of plants grown in liquid soil extract (23 genes all belonging to the same cluster, ABIE53_000074- ABIE53_000114, but not in the other substrates (**Fig. 5a**). Those 23 genes are related to flagellum synthesis (**Fig. 5b**, **Fig. 5c**) and swimming motility (**Fig. 5c, Table S10**), as we could confirm. They also have a positive average fitness for many of the *in vitro* assays we performed (**Table S10**). Interestingly, the flagellar hook-associated protein FlgK (ABIE53_000091) and the c-di-GMP-binding flagellar brake protein YcgR (ABIE53_000092) (**Fig. 5b**) have very different fitness values from the rest of the cluster, not only in plant but also in several *in vitro* conditions (**Table S10**), suggesting that these genes play regulatory roles rather than just structural components of the flagella.

With the purpose of validating the effect of plant growth substrate on plant colonization fitness of strain OAS925, we generated an archived collection of transposon mutants and selected individual mutants to test in competition with the wild-type strain in 1:1 proportion. We selected individual transposon mutants of genes ABIE53_000305 (general secretion pathway protein L), ABIE53_001142 (hypothetical protein), ABIE53_002867 (2-isopropylmalate synthase), ABIE53_003398 (zinc/manganese transport system substrate-binding protein), ABIE53_003440 (cyclic pyranopterin phosphate synthase, related to molybdenum cofactor biosynthesis), ABIE53_004277 (3- oxoadipate CoA-transferase alpha subunit), and ABIE53_004744 (methyl-accepting chemotaxis protein), and evaluated their *Brachypodium* colonization ability in two plant growth substrates in which they had shown contrasting fitness values (**Table S10**) and in R2A media as a control. For all the mutants we could validate the difference of fitness in two selected substrates (**Fig. 6**) and an absence of phenotype in R2A rich media (data not shown). For example, the ABIE53_003398 mutant colonized poorly in liquid soil extract relative to 05X MS agar, in agreement with the RB-TnSeq data. Overall, the data confirmed the results obtained in our RB-TnSeq screening and the difference of fitness observed in the competition assays was consistent with the difference predicted by RB- TnSeq (**Fig. 6b**). Together, these validation efforts support our conclusion that plant growth substrate influences bacterial colonization fitness.

**Figure 6.**
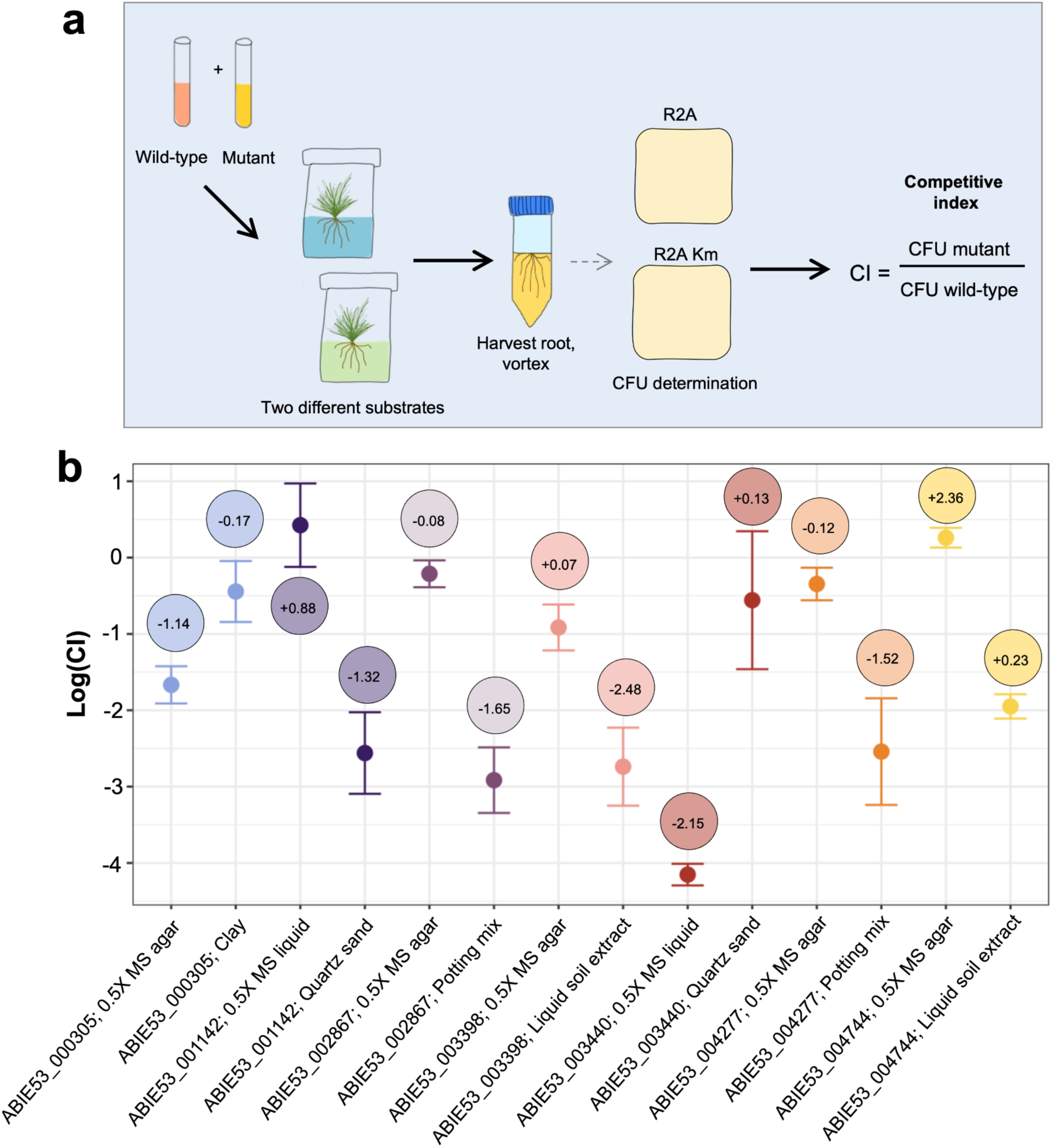
Validation of RB-TnSeq results with individual transposon mutants of *Paraburkholderia graminis* OAS925. **a**) Overview of the plant competition assays between each mutant and the wild-type strain OAS925. **b**) Plot shows competitive index (CI) values for each mutant in competition against the OAS925 wild-type strain in *Brachypodium distachyon* Bd21-3 grown in two selected plant growth substrates. Mutants in genes ABIE53_003440 (cyclic pyranopterin phosphate synthase, related to molybdenum cofactor biosynthesis), ABIE53_003398 (zinc/manganese transport system substrate-binding protein), ABIE53_004744 (methyl-accepting chemotaxis protein), ABIE53_001142 (hypothetical protein), ABIE53_004277 (3-oxoadipate CoA-transferase alpha subunit), ABIE53_002867 (2-isopropylmalate synthase) and ABIE53_000305 (general secretion pathway protein L) are shown. For illustration purposes, log transformation is shown. For comparison, average fitness value from RB-TnSeq assays is represented in circles (data obtained from Table S8). CI values were calculated using at least 3 replicates per condition as colony forming units (CFU) of mutant/CFU wild-type. CI &lt; 1 means the mutant is less fit than the wild- type, CI > 1 means the mutant is more fit than the wild-type.

### TnSeq across many in vitro conditions reveals functions of unannotated rhizosphere colonization genes in OAS925

Genome-wide mutant fitness data across many conditions is useful for defining functions for genes currently annotated with vague or hypothetical functions (Price *et al*., 2018). From our OAS95 fitness dataset, we could link genotype to phenotype for 3,758 genes (|fitness| ≥ 1; **Fig. S6a**); or 1,807 genes if considering both |fitness| ≥ 1, |t| ≥ 3 (**Fig. S6b**). This allowed us to hypothesize about the role of unannotated rhizosphere colonization genes (hypothetical function, COG class S).

For instance, ABIE53_003575 (detrimental for colonization in 0.5X MS agar; unannotated homologues in *Burkholderia* and *Paraburkholderia* species) is also detrimental (fitness ≥ 1, t ≥ 3) for metabolism of D-serine, D-ribose, 2-deoxy-D-ribose, gluconic, carnitine and asparagine; but it is important (fitness ≤ -1, t ≤ -3) for growth with 4-hydroxybenzoic acid (**Table S10**). ABIE53_003575 has a highly correlated pattern of phenotypes with the adjacent ABIE_003574 (annotated as a low-complexity acidic protein), suggesting that these two genes function together. ABIE53_005492 (important for plants in liquid soil extract; annotated as putative tetrahydromethanopterin-linked C1 transfer pathway protein) is also important for arabinoxylan metabolism (**Table S10**). ABIE53_000326 (annotated as a hypothetical protein; has homology with BP1026B_I0091 surface attachment protein Sap1 from *Burkholderia pseudomallei* MSHR146 (Sun *et al*., 2023)), which is detrimental for colonization of plants in 0.5X MS agar, is also detrimental for metabolism of D-lactate. Despite the findings above, there were some cases of rhizosphere genes (e.g. putative membrane protein ABIE53_003209 and hypothetical protein ABIE53_001726; **Table 1, Table S11**) for which we could not link *in vitro* phenotypes that explained their potential function.

Apart from better understanding the role of unannotated plant rhizosphere colonization genes, conducting numerous *in vitro* assays also allowed us to identify a specific phenotype for 86 unannotated or poorly characterized non-rhizosphere genes in the genome of *P. graminis* OAS925 (**Table S13**), and for some of them we could propose a specific role within a process. Genes could be associated with a specific condition because the gene affected had fitness only in that unique condition (or a small subset of conditions) (|fitness| ≥ 1, |t| ≥ 3). For instance ABIE53_003311 is required for tolerance to limonene (a plant-related monocyclic monoterpene), ABIE53_000770 is important for D-lactate catabolism, ABIE53_000853 is involved in the metabolism of azelaic acid, and ABIE53_005694 (encoding a LysR-class transcriptional regulator) is important for growth with L-rhamnose (**Table S13**). ABIE53_005694 is in the center of a gene cluster (ABIE53_005700-ABIE53_005690) which is annotated as involved in L-rhamnose transport and catabolism, and hence is likely a regulator specific for this compound.

### 3.4. Bacterial fitness substrate dependency is observed in other plant-related species

To determine if type of substrate impacts colonization fitness in other bacteria, we investigated three additional plant-associated bacteria, the beneficial strain *P. phytofirmans* PsJN and the switchgrass isolates *Variovorax* sp. OAS795 and *Rhizobium* sp. OAE497, and we conducted experiments with *Brachypodium* plants grown in two different types of substrates. According to confocal microscopy, strains PsJN and OAE497 colonize *B. distachyon* (**Fig. 7a**) in a similar manner to OAS925 (**Fig. 2**). For OAE497 we could observe putative endophytic (i.e. inside root cells) growth (**Fig. 7a**). In contrast, OAS795 seems to colonize less efficiently and to have a preference for the plant cells liberated in the root cap (**Fig. 7a**).

**Figure 7.**
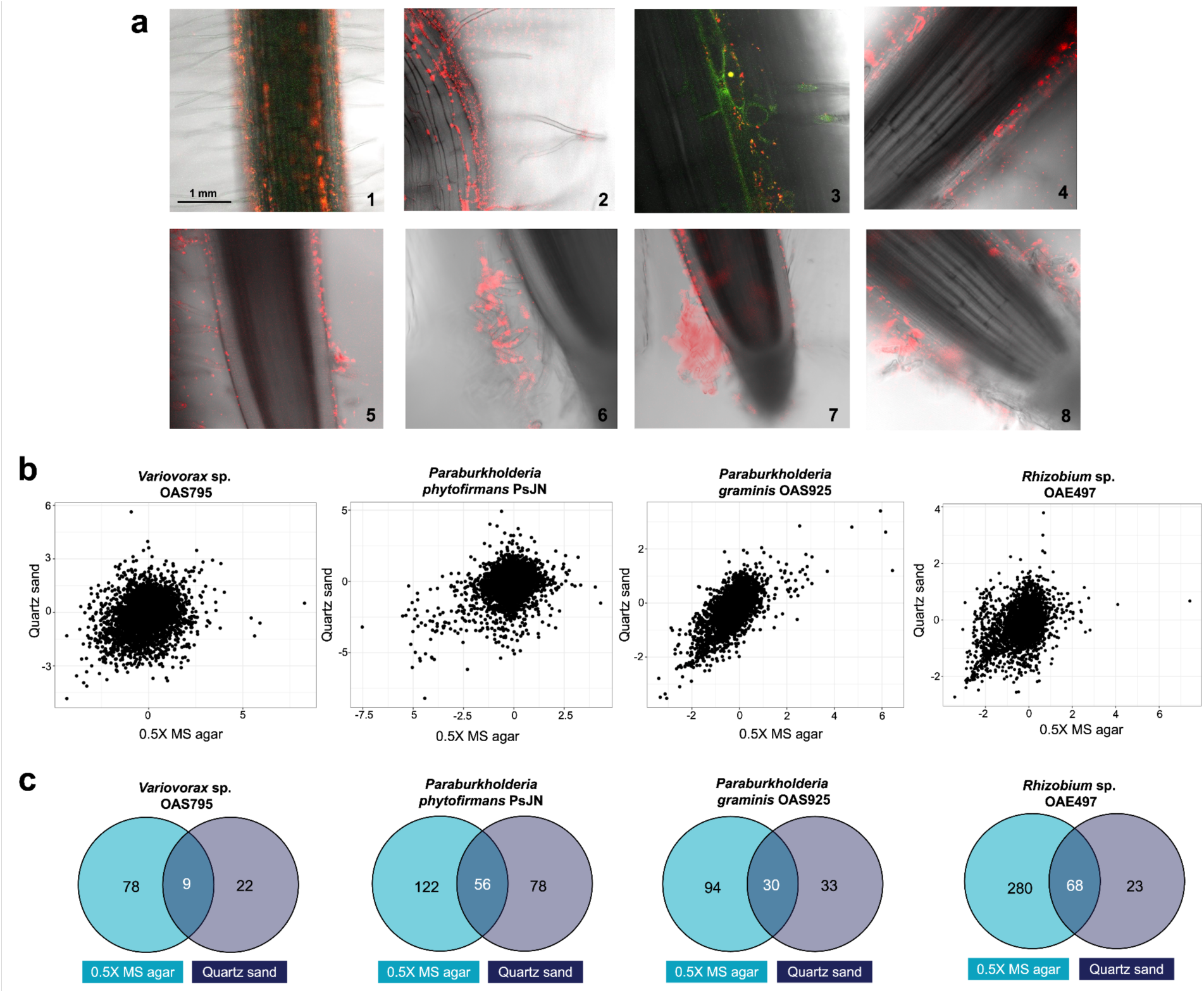
Influence of plant growth substrate on colonization bacterial fitness of *Variovorax* sp. OAS795, *Rhizobium* sp. OAE497 *Paraburkholderia phytofirmans* PsJN, and *P. graminis* OAS925. a) Confocal microscopy observation of *Brachypodium distachyon* Bd21-3 roots colonized by red fluorescent protein (RFP) tagged derivative strains of PsJN (1), OAE497 (2, 3) and OAS795 (4, 5, 6, 7, 8). Colonization was observed 7 days after inoculation at 20X and 40X amplification using a Zeiss LSM 710 confocal microscope system. Overlay of fluorescence and bright-field microscopy is shown. b) Correlation of fitness colonization data of *B. distachyon* Bd21-3 plants grown in 0.5X MS agar versus quartz sand. Plots were generated using average fitness values for replicate experiments. c) Venn diagrams showing the number of unique and shared genes meeting the |fitness| ≥ 1 and |t| ≥ 3 criteria in 0.5X MS agar and quartz sand in each of the bacterial species.

We used barcoded mutant libraries of PsJN, OAS795, and OAE497. The PsJN RB-TnSeq library (named BFirm_ML3) was already described (Price *et al*., 2018), and generated fitness data for 5,428 genes (74.% of the genome). For OAS795, mutagenesis and subsequent TnSeq resulted in a pool of 78,640 individual mutant strains (library *Variovorax*_OAS795_ML2) with mapped insertions and unique barcodes at 41,793 different locations distributed across the genome at an average rate of 1 insertion every 146 bp. From 5,899 genes in its genome, we generated fitness data for 4,532 genes (76.8% of the genome). For OAE497, mutagenesis and subsequent TnSeq resulted in a pool of 557,956 individual mutant strains (library *Rhizobium*_OAE497_ML4) with mapped insertions and unique barcodes at 139,136 different locations distributed across the genome at an average rate of 1 insertion every 6 bp. From 6,194 genes in its genome, we generated fitness data for 5,152 genes (83.2% of the genome). In all the mutant libraries, mapped insertions had locations distributed across the genome (**Fig. S7**).

Colonization fitness experiments were conducted in plants grown in two types of substrate, quartz sand and 0.5X MS agar. These substrates were chosen due to 1) their frequent use in gnotobiotic set ups in plant research and 2) the differences observed among them in root and shoot weight (**Fig. 3a**, **Fig. 3b**). When comparing the plant colonization fitness data in quartz sand versus 0.5X MS agar for OAS795 (**Table S14**), PsJN (**Table S15**), OAE497 (**Table S16**), and OAS925 (**Table S8**) we observed low correlation (**Fig. 7b**). The higher correlation observed for OAS925 (R = 0.65) compared to the other three strains (OAS795 R = 0.23, PsJN R = 0.32, OAE497 R = 0.41) may be explained by the higher number of replicates for OAS925 (OAS925 n = 9; OAS795 n = 2- 5; PsJN n = 1-5; OAE497 n = 2) (**Table S7**, **Table S14**, **Table S15**). Furthermore, the genes identified as colonization factors (|fitness| ≥ 1 and |t| ≥ 3) tended to be unique to each substrate (**Fig. 7c**). Although it is not the main scope of this work to discuss this, we observed some interesting substrate-dependent phenotypes in each of the strains. For instance, in *Rhizobium* sp. OAE497 we found that a large cluster involved in flagellar synthesis (ABIE40_RS02910-ABIE40_RS03125) was important for colonization of plants grown in 0.5X MS agar but detrimental for plants grown in quartz sand (**Table S16**). This cluster was also detrimental for metabolism of L-asparagine (Robin Herbert, personal communication), which is more abundant in the quartz sand condition than in 0.5X MS agar (**Fig. 3**). Flagellar motility is related to efficient host colonization, dispersion, and attachment (Knights *et al*., 2021) and has recently been shown to be linked to carbon dynamics in soil (Ramoneda *et al*., 2024), and therefore it could be proposed as a good target to engineer in beneficial strains and improve their plant colonization abilities. However, our results on the influence of substrate type on flagellar gene fitness for the strains OAE497 (**Table S16**) and OAS925 (**Fig. 5, Table S10**) point to different bacterial lifestyles across plant growth substrates, and they show that mutating flagellar genes does not necessarily give an advantage on all plant growth substrates. Overall, these results strengthen our recommendations on using different growth substrates when assessing plant-bacteria interactions.

In order to facilitate the comparison of the influence of plant growth substrate on fitness genes across different species, we used orthologue data and focused only on the three species of the order Burkholderiales (*P. graminis* OAS925, *P. phytofirmans* PsJN, *Variovorax* sp. OAS795). We found core fitness genes shared in all three species that are important for plant colonization in both of the substrates. One example is the cytochrome c oxidase assembly protein ABIE53_000682 (orthologue in PsJN is BPHYT_RS02685, orthologue in OAS795 is ABID97_RS23920), that also has a negative fitness for 2-deoxy-D-ribose metabolism in the three species (**Table S7**, **Table S14**, **Table S15, Table S17** and (Wetmore *et al*., 2015; Price *et al*., 2019)). Another example is the glutamate synthase ABIE53_003734 (orthologue in PsJN is BPHYT_RS17855, orthologue in OAS795 is ABID97_RS23225) (**Table S17**), which also has negative fitness for 2-deoxy-D-ribose, D-xylose, D-alanine, D-fructose and L-malic acid in the three species. Interestingly, both ABIE53_000682 and ABIE53_003734 genes are part of the OAS925 core colonization genes needed across different plant growth substrates and are also involved in colonization of *Medicago* plants (**Table S12**); and their homologues in OAS795 and PsJN are also needed across the different substrates tested (**Table S14**, **Table S15**).

Apart from the above core colonization genes, we also identified plant fitness genes that are influenced by substrate type in multiple bacteria. Briefly, we looked for PsJN and/or OAS795 orthologues in the 382 OAS925 colonization genes, and calculated the |fitness difference in the two substrates| (**Table S17**). We found that some of the soil-influenced genes (|fitness difference in the two substrates| ≥ 1) in the two closely-related OAS925 and PsJN strains did not have a different pattern across substrates in *Variovorax* sp. OAS795. Examples are genes involved in magnesium/cobalt transport (ABIE53_003483), glycolysis/gluconeogenesis (ABIE53_002485) and molybdenum biosynthesis (ABIE53_003060-ABIE53_003062) (**Table S17**). Some of the genes in this last group also have a strong negative fitness in OAS925 for growth with adenosine as the sole carbon source (**Table S8**). We found few examples of orthologous genes in the three species that are impacted by the type of substrate. Examples of genes that are more important for colonization of plants in one substrate than the other include: ABIE53_000656 (gamma-glutamyl cycle/biosynthesis of glutathione) and ABIE53_003282-ABIE53_003284 (amino acid transport genes; needed for fitness *in vitro* with amino acids such as L-proline and L-asparagine) (**Table S8**, **Table S14**, **Table S15** and (Wetmore *et al*., 2015)). Interestingly, the main substrate-binding component of this ABIE53_003282-ABIE53_003284 transporter is encoded elsewhere in the genome (ABIE53_000461) and has the same phenotype in plants and *in vitro* as the transporter genes. Globally, these results reinforce that there is a strong effect of plant growth substrate on plant colonization fitness in diverse bacteria, and they highlight the importance of considering different substrates when studying plant-bacteria interactions.

## 4. Discussion

Plant roots are colonized by a diverse microbiome, which is recruited from surrounding soil communities and modulated by host plant immunity and root metabolites (Jones & Dangl, 2006; Bulgarelli *et al*., 2012). Understanding the molecular, genetic, and ecological mechanisms that govern plant-bacteria interactions is pivotal for advancing sustainable agriculture and ecosystem health. Among the different factors that affect soil and rhizosphere bacterial communities, substrate type is one of the most disregarded, although 16S rRNA sequencing data has defined plant growth substrate as one of the main determinants of the composition of bacterial communities in soils (Girvan *et al*., 2003). Soil chemistry and particle size shape root morphology, root exudate composition, and metabolite availability (Neumann *et al*., 2014; Swenson *et al*., 2015; Sasse *et al*., 2020). In the same way that soil influences the composition of root exudates, these can influence the physical properties of soil (e.g. pH, water retention, or aggregate stability), as well as impact the soil microbial community structure. Different soil properties can influence bacterial motility and biofilm formation (Ma *et al*., 2017; Yang & Van Elsas, 2018), due to the pore network characteristics, and the availability of bacteria-bacteria and plant-bacterial communication signals (Del Valle *et al*., 2020, 2022). Understanding the effect that soil has on rhizospheric microorganisms remains challenging because parameters vary simultaneously and uncontrollably when different types of soils/substrates are compared. Additionally, it is not easy to disentangle the effects of soil physical characteristics from the effect of soil-dependent differentially-produced root exudate compounds on bacteria, therefore complicating mechanistic determination of any individual soil parameter’s effect on microbial behaviors.

In order to evaluate the impact of plant growth substrate in plant colonization fitness genes we constructed an RB-TnSeq mutant library in *P. graminis* OAS925, a strain that is able to compete with other soil microorganisms and efficiently colonize plant roots (Lin *et al*., 2023). As a model plant we chose *B. distachyon*, a grass crop in which the rhizosphere is dominated by bacteria from the order Burkholderiales (Kawasaki *et al*., 2016). We performed rhizosphere colonization assays in *B. distachyon* plants grown gnotobiotically in six different growth substrates, and compared the fitness of *Paraburkholderia*_OAS925_ML2 across them. The growth substrates were chosen due to their frequent use in gnotobiotic experiments: 0.5X MS liquid, 0.5X MS agar, clay, quartz sand, potting mix, and liquid soil extract. In total, we identified 382 genes to be involved in *B. distachyon* rhizosphere colonization in at least one plant growth substrate.

Carbohydrates, amino acids, organic acids, and phenolic compounds in root exudates of *B. distachyon* have been shown to feed soil bacteria. In *Pseudomonas* species, for instance, they alter the expression of diverse metabolic, transport, regulatory, and stress genes (Mavrodi *et al*., 2021). In *Bacillus* species they are chemoattractants and stimulate biofilm production (Sharma *et al*., 2020). Here, we show that *P. graminis* OAS925 can grow on different compounds that are known to be present in *B. distachyon* exudates, such as histamine, xanthine, histidine, threonine, arginine, asparagine, leucine, proline, azelaic acid, and salicylic acid (Sasse *et al*., 2019; Sharma *et al*., 2020). We found that all of these compounds are differentially abundant across plant growth substrates, which agrees with previous analyses on *B. distachyon* plants grown on diverse substrates (Sasse *et al*., 2019, 2020).

Contrary to *in vitro* fitness assays, where a link between genotype and phenotype can more easily be established, genome-wide assessment of plant colonization considerably exceeds *in vitro* assays in terms of complexity. The dynamic and heterogeneous nature of plant colonization makes it challenging to untangle the role of each of the 382 bacterial fitness genes involved in rhizosphere colonization in *P. graminis* OAS925. We approached this using 110 *in vitro* assays that mimic different conditions in the rhizosphere environment to further characterize the physiological functions of the genes in more detail. The latter included metabolism of 76 carbon and 6 nitrogen sources commonly found in root exudates, tolerance to 20 different stress conditions frequently found in the plant environment, motility, and chemotaxis to plant roots. We also tested the mutant library on *A. thaliana* and *M. truncatula* plants grown in quartz sand and potting mix, respectively.

Although most of the 382 rhizosphere colonization genes that we identified in *P. graminis* OAS925 only had a fitness in one or a few of the plant growth substrates tested, a core set of 34 genes were involved in plant colonization across all the plant growth substrates, which makes them attractive targets for manipulation of the colonization abilities of strain OAS925. This represents <14% of the total number of genes identified in plants grown in liquid soil extract, reflecting the risk of considering only one substrate when assessing bacterial plant colonization fitness. Some of these 34 core colonization genes have previously been described in plant bacteria as involved in colonization; for instance, ABIE53_003556 (galactonate metabolism) (Cole *et al*., 2017) and ABIE53_000413-ABIE53_000414 and ABIE53_004817 (methionine synthesis) (Sivakumar *et al*., 2019; Su *et al*., 2021), which reflects that aspects of colonization mechanisms are shared across bacteria, despite their taxonomic diversity.

We identified 348 genes that only showed fitness in one or a few substrates, and thus are colonization factors that are influenced by the environment. Several genes even had opposite-direction fitness values depending on the type of substrate considered. For instance, genes in the cluster ABIE53_001141-ABIE53_001160 (coding for glycosyltransferases, lipopolysaccharide transport systems and hypothetical proteins) are detrimental for colonization of plants in 0.5X MS liquid and liquid soil extract, but are needed for colonization of plants grown in 0.5X MS agar, quartz sand and potting mix. Examples of genetic determinants that show a significant fitness effect only in one or a few substrates include: ABIE53_000074-ABIE53_000114 (flagellar synthesis), ABIE53_004752-ABIE53_004749 (spermidine/putrescine transport), ABIE53_000295- ABIE53_000307 (general secretion pathway), ABIE53_004744 (methyl-accepting chemotaxis protein) and ABIE53_003400-ABIE53_003398 (zinc/manganese transport). Many of these genes were previously related to efficient plant colonization. For several of them, we could validate the observed substrate dependency on the colonization fitness using individual mutants in competition with the wild-type strain.

The different plant growth substrates that we used in this study have diverse metabolite composition, density, porosity, and aeration; and hence establishing a direct link between soil relevant parameters, root exudate composition, and rhizosphere colonization fitness remains challenging. One example of lack of correlation is observed for ABIE53_002392, which encodes a predicted OHCU decarboxylase (homology with uricase), and putatively acts in the metabolism of purines such as adenosine and xanthine. We confirmed the importance of this gene for growth of OAS925 on adenosine and xanthine as nutrients, and detect it in our colonization assays in plants grown in 0.5X MS liquid (in which adenosine is most abundant) and liquid soil extract (where xanthine is most abundant), but we also detect it in 0.5X MS agar where these two purines are lower in abundance. Because rhizosphere metabolites are a mixture of many compounds, disrupting the utilization of one nutrient might not lead to a measurable loss of fitness. Similarly, genes involved in molybdenum metabolism (ABIE53_002030, ABIE53_003060-ABIE53_003062, ABIE53_003118 and ABIE53_003440) are important for colonization of plants in 0.5X MS agar but not in quartz sand in both OAS925 and PsJN. *In vitro*, some of these same molybdenum cofactor biosynthesis genes are needed for growth with adenosine. However, our metabolomic analyses show that adenosine is more abundant in the rhizosphere of plants grown in quartz sand than in 0.5X MS agar. Interestingly, molybdenum cofactor synthesis genes in *Burkholderia* species have been related to biofilm formation (Andreae *et al*., 2014). Our different fitness results for molybdenum cofactor synthesis genes between the aforementioned plant growth substrates could therefore reflect an impact in a biofilm phenotype instead of a difference in nutrient utilization, suggesting that that rhizosphere metabolites are not the sole drivers of bacterial fitness in the plant environment.

Despite the challenge of linking plant growth substrate, root exudate composition, and bacterial plant fitness, we could identify a more direct connection in some cases. For instance, ABIE53_004277 is only important for rhizosphere colonization of *B. distachyon* plants grown in potting mix. This gene codes for 3-oxoadipate CoA-transferase alpha subunit, which is involved in the catabolism of a variety of aromatic compounds such as benzenoid derivatives. In our metabolomic analysis, we found that dihydroxybenzoic acids (benzenoid derivatives) are more abundant in the potting mix than in any other substrate, which therefore correlates with our finding. With our *in vitro* fitness data we could confirm that ABIE53_004277 is important in the metabolism of 4-hydroxybenzoic acid.

Altogether, our results show that RB-TnSeq is a useful tool to find functions for poorly annotated genes and discover novel plant colonization genes. We demonstrate that RB-TnSeq can define colonization differences across conditions and that bacterial fitness for plant colonization is strongly influenced by growth substrate. When doing TnSeq-like studies in root and rhizosphere systems, we recommend using different types of substrates and a high number of replicates, especially when using growth substrates with high variability, such as potting mix. The results of our research highlight the importance of considering different growth substrates in plant-bacteria interaction studies, especially when selecting fitness genes for targeted manipulation of the colonization capabilities of plant-beneficial or pathogenic bacteria.

## Acknowledgments

The authors thank John Vogel and Mingqin Shao for providing Bd21-3 seeds and clay, Mitchel Thompson for sharing plasmid pGingerBK-J23100, Peter Kim for providing Imaging EcoFABs, Hans Carlson for sharing carbon sources and assisting with the arrayed library creation, Alexey Kazakov for assisting with gene orthologue data, and the QB3 Genomics Center at UC Berkeley for sequencing.

## **7.** Author contributions

MT and AMD designed and coordinated the study. MT constructed mutant libraries, performed fitness assays and validations *in planta*. AK and AMD assembled the arrayed collection of individual OAS925 transposon mutants. AK and KZ collected, processed and analyzed rhizosphere metabolites. SMK coordinated LC-MS/MS analysis and finalized metabolite annotations. MNP processed raw sequencing data and calculated gene fitness scores. TRN and AMD obtained funding. MT analyzed data, prepared figures and tables, and wrote the manuscript. All authors reviewed the final version of the manuscript.

## Supporting Information

**Note S1.** *Brachypodium distachyon* seed sterilization.

**Note S2.** Construction of a barcoded transposon library in *Paraburkholderia graminis* OAS925.

**Note S3.** Preparation of plant growth substrates.

**Note S4.** Collection and identification of rhizosphere metabolites.

**Note S5.** RB-TnSeq *in vitro* growth fitness assays.

**Note S6**. DNA isolation, library preparation and sequencing.

**Note S7**. Construction of RB-TnSeq libraries in *Variovorax* sp. OAS795 and *Rhizobium* sp. OAE497.

**Table S1**. Arrayed collection of *Paraburkholderia graminis* OAS925 transposon mutants.

**Table S2**. In-house library of authentic reference standards used for rhizosphere metabolite identification.

**Table S3**. Summary of fitness assays conducted with *Paraburkholderia*_OASA925_ML2.

**Table S4.** Summary of fitness assays conducted with *Variovorax*_OAS795_ML2, *Rhizobium*_OAE497_ML4 and BFirm_ML3.

**Table S5**. Rhizosphere metabolites in *Brachypodium distachyon* Bd21-3 plants grown in different types of substrates. Peak heights are shown.

**Table S6**. *Paraburkholderia*_OASA925_ML2 gene information, including scaffold, begin, end, strand, GC content of the gene’s sequence and nTA (number of TA dinucleotides in the gene’s sequence; the *mariner* transposase usually inserts at TA).

**Table S7**. *Paraburkholderia*_OASA925_ML2 gene fitness and t values across all conditions tested, and list of assays that are available at the Fitness Browser website (https://fit.genomics.lbl.gov).

**Table S8**. *Paraburkholderia*_OASA925_ML2 average fitness and t values across all conditions tested.

**Table S9**. *Paraburkholderia*_OASA925_ML2 genes with a fitness defect or advantage (|fitness| ≥ 0.75 and

|t| ≥ 4) in the outgrowth controls.

**Table S10**. *Paraburkholderia*_OASA925_ML2 genes involved in *Brachypodium distachyon* Bd21-3 rhizosphere colonization across one or more types of substrates (|fitness| ≥ 1 and |t| ≥ 3 in at least one substrate type). Fitness and t values of other plant and *in vitro* assays are also shown.

**Table S11**. *Paraburkholderia*_OASA925_ML2 genes involved uniquely in *Brachypodium distachyon* Bd21- 3 rhizosphere colonization across one or more types of substrates (|fitness| ≥ 1 and |t| ≥ 3 in at least one substrate type). Fitness and t values of other plant and *in vitro* assays are also shown.

**Table S12**. *Paraburkholderia*_OASA925_ML2 genes involved in *Brachypodium distachyon* Bd21-3 rhizosphere colonization across all types of substrates (|fitness| ≥ 1 in all plant growth substrates, and |t| ≥ 3 in at least one substrate type). Fitness and t values of other plant and *in vitro* assays are also shown.

**Table S13**. Unannotated genes in *Paraburkholderia*_OASA925 with one unique phenotype *in vitro* (|fitness|

≥ 1 and |t| ≥ 3). Fitness and t values of plant and *in vitro* assays are also shown.

**Table S14**. *Variovorax*_OAS795_ML2 gene fitness and t values, and list of assays that are available at the Fitness Browser website (https://fit.genomics.lbl.gov).

**Table S15**. BFirm_ML3 (*Paraburkholderia phytofirmans* PsJN ML3) gene fitness and t values, and list of assays that are available at the Fitness Browser website (https://fit.genomics.lbl.gov).

**Table S16**. *Rhizobium*_OAE497_ML4 gene fitness and t values, and list of assays that are available at the Fitness Browser website (https://fit.genomics.lbl.gov).

**Table S17**. Comparison of substrate influence on *Brachypodium distachyon* average fitness colonization data for *Paraburkholderia graminis* OAS925, *P. phytofirmans* PsJN and *Variovorax* sp. OAS795. Average fitness values are obtained from averaging replicates in Table S8, Table S14 and Table S15.

---

Figure S1. **a**) Overview of *Brachypodium distachyon* Bd21-3, *Medicago truncatula* A17 and *Arabidopsis thaliana* Col-0 plants grown in different types of substrate: 0.5X MS agar, 0.5X MS liquid, clay, liquid soil extract, potting mix, and quartz sand. **b**) Different root morphology (hair roots) observed in *Brachypodium* plants grown in 0.5X MS liquid and liquid soil extract.

Figure S2. “Chemotaxis to plant root” assay. *Paraburkholderia*_OAS925_ML2 was inoculated at ∼1 cm (black dot, where the bubbles appear) from 21-old-day *Brachypodium distachyon* Bd21-3 plants that were transferred from pots with 0.5X MS with 0.3% w/v agar onto plates with 0.5X MS with 0.25% w/v agar the day before inoculation. Twenty four h after inoculation, outer samples (arrows) with motile cells were harvested.

Figure S3. Average relative abundance of compounds across different plant growth substrates; in some cases compound names in panels have been shortened for illustration purposes, see Table S5. Different letters indicate statistically significant differences (two-way analysis of variance (ANOVA) with Tukey’s post hoc test; *P* ≤ 0.05).

Figure S4.Heatmap of average fitness values for the 34 core rhizosphere colonization genes across *Brachypodium* plants grown in different plant growth substrates and different *in vitro* conditions. Gene fitness values are bounded from -2 to 2.

Figure S5. Influence of plant growth substrate on *Paraburkholderia graminis* OAS925 bacterial fitness for plant colonization. **a**) Total number of genes with negative or positive fitness values in the different substrates (|fitness| ≥ 1 and |t| ≥ 3). **b**) Shared and unique fitness colonization genes (|fitness| ≥ 1 and |t| ≥ 3) across plant growth substrates. Gray boxes show the number of shared genes across substrates; dashed empty boxes mean genes are not required for that type of substrate. For illustration purposes, numbers (n ≤ 6) corresponding to genes shared in a few substrates only are not shown in the top figure. **c**) Heatmap of fitness values across the different plant growth substrates for a subset of genes; genes with large differences in fitness between two different substrates (calculated as the |maximum fitness - minimum fitness|) are shown. Gene fitness values for each replicate experiment are plotted separately. Gene fitness values are bounded from -2 to 2.

Figure S6. Number of genes with a fitness phenotype after conducting experiments in a given number of conditions. **a**) Criteria is |fitness| ≥ 1. **b**) Criteria is |fitness| ≥ 1 and |t| ≥ 3.

Figure S7. Histograms of mapped insertions for *Variovorax*_OAS795_ML2, BFirm_ML3, *Rhizobium*_OAE497_ML4 and *Paraburkholderia*_OAS925_ML2.

