## Supplementary Material for "Bacterial fitness for plant colonization is influenced by plant growth substrate"

*Adam M. Deutschbauer 0000-0003-2728-7622*

### Notes:

**Note S1. *Brachypodium distachyon* seed sterilization.**

*B. distachyon* Bd21-3 seeds were dehusked and sterilized in 70% v/v ethanol for 30 s, and in 6% v/v NaOCl for 5 min, followed by five wash steps in sterile water. Seeds were stratified in the dark for 2 to 3 days at 4ºC. After stratification, seeds were germinated on 1% w/v water-agar plates in a 130 μmol/$m^2$ $s^{-1}$ 16-h light/8-h dark regime at 26°C for three days.

**Note S2. Construction of RB-TnSeq library in *Paraburkholderia graminis* OAS925.**

We grew 10 mL of wild-type OAS925 in R2A overnight at 30ºC. The next morning, we recovered a 2 mL freezer stock of strain AMD290 in 50 mL LB supplemented with 50 μg/mL Cb and 300 μM DAP at 37ºC. When the OD600 of the *Escherichia coli* donor strain reached 1, we harvested 20 OD600 units of the culture and washed the cells three times with fresh LB supplemented with DAP. Then, 20 OD600 units of wild-type OAS925 cells were harvested, mixed with the washed donor cells, centrifuged, and resuspended in a final volume of 0.5 mL with LB supplemented with DAP. The resuspension was spotted onto 0.45-μm membrane filters (Millipore, United States) and incubated overnight on LB agar plates supplemented with DAP at 30ºC. The next day, the conjugation mixture was scraped from the membrane, resuspended in 10 mL R2A with 10 μg/mL Km and plated at different dilutions on R2A plates supplemented with 10 μg/mL Km. Plates were incubated at 30ºC for 48 h to let visible colonies develop. We then pooled ~400,000 colonies and grew the library in liquid R2A supplemented with 10 μg/mL Km for 2 population doublings. We then added glycerol to a final volume of 15% v/v, made multiple 1-mL -80°C freezer stocks (~$10^8$ cells/mL) of the final library for subsequent experiments, and collected cell pellets to extract genomic DNA for TnSeq mapping.

**Note S3. Preparation of plant growth substrates.**

Quartz sand (50-70 mesh particle size; Sigma-Aldrich, United States), clay (porous ceramic clay, Greens grade; Profile Products, United States), 0.5X MS liquid and 0.5X MS with 0.3% w/v agar were sterilized by autoclaving for 20 min at 121°C. Potting mix

(Miracle Gro, United States) was sterilized by autoclaving twice for 30 min at 121°C, with a 24 h-incubation at 30ºC between autoclave cycles. Liquid soil extract was prepared by mixing 100 g of potting mix (Miracle Gro, United States) in 1 L of water by gentle shaking, followed by 16 h of incubation at 4°C with continuous stirring, and filtration through Miracloth (Millipore, United States) and then a 0.22-μm membrane filter for sterilization. Dry substrates (quartz sand, clay, potting mix) were moistened with 35 mL of 0.1X MS before transferring the seedlings into the plant culture boxes.

**Note S4. Collection and identification of rhizosphere metabolites.**

Four weeks after transplanting the seedlings into the substrates, plants were harvested with tweezers. For liquid substrate conditions, 18 mL of medium were collected, filtered through a 0.45 μM filter (Acrodisc filter with Supor membrane; Pall Life Sciences), and stored at -80ºC until analysis. For the solid substrate conditions, a sterile spatula was used to remove bulk soil from the roots, which were then cut from shoot using a scalpel and transferred into 50-mL Falcon tubes with 18 mL of fresh MS medium. The roots were then vortexed for 10 s at maximum speed to collect and remove the loosely bound substrate. Using tweezers, roots were then removed and the substrate/medium mix was shaken for 1 h at 180 rpm at 4ºC to extract water soluble metabolites from the substrate particles. The solution was centrifuged at 6,000 *g* for 10 min in order to pellet the substrate, and then the supernatant was collected and filtered through a 0.45 μM filter (Acrodisc filter with Supor membrane; Pall Life Sciences); filtrate was collected and stored at -80ºC for further analysis. Four replicates were used per condition.

Ten mL of frozen samples for LC-MS analysis were lyophilized; and 1 mL of 100% ice-cold LC-MS grade methanol (Honeywell Burdick & Jackson, United States) was added. Samples were vortexed for 10 s, sonicated for 30 min on ice and incubated at 4ºC overnight. Samples were centrifuged at 10,000 *g* for 10 min at 4ºC. Supernatant was dried in a Savant SpeedVac SPD111V (Thermo Scientific, United States) for 3 h. Dried samples were sonicated for 15 min using an ultrasonic bath with 300 uL of 100% ice-cold LC-MS-grade methanol with internal standards (Table S2). Resuspended samples were filtered with 0.22 μm microcentrifuge PVDF filters (Merck Millipore, United States) and 250 μL aliquots were transferred to vials for LC-MS analysis (Baker et al., 2024). Metabolites were separated using a HILIC-Z column on an Agilent 1290 LC stack with MS and MS/MS data collected using a Thermo Q Exactive Hybrid Orbitrap Mass Spectrometer (Thermo Scientific, United States).

**Note S5. RB-TnSeq *in vitro* growth fitness assays.**

Carbon source assays were conducted in RCH2_defined_noCarbon supplemented with different carbon sources (**Table S3**). Washed cells were inoculated in the different media (starting OD600 = 0.02) and incubated at 30ºC in a multitron shaker at 700 rpm. Nitrogen source assays (**Table S3**) were conducted as the carbon source assays above but using RCH2_defined_noNitrogen instead of RCH2_defined_noCarbon. Stress assays were conducted in R2A media using compounds (**Table S3**) at inhibitory, but sub-lethal concentrations. After 24, 48, 72 h, or 96 h, cultures with an OD lower than the no stress controls were centrifuged. Pellets were kept frozen at -20ºC until DNA purification. Unless noted otherwise, chemical compounds were purchased from Sigma-Aldrich (United States).

**Note S6. DNA isolation, library preparation and sequencing.**

Purified genomic DNA was measured on a nanodrop device and ~200 ng of RNA-free DNA was used as a template for DNA barcode PCR amplification using previously described PCR conditions with Q5 DNA polymerase with Q5 high GC enhancer (New England Biolabs, United States). Briefly, DNA was used as a template in a PCR using primers flanking the transposon barcode region, each containing an Illumina adapter and multiplexing index sequence. The BarSeq primers contained unique indexes on both primers to identify and remove instances of index hopping. BarSeq PCR amplicons were sequenced on an Illumina instrument (either HiSeq2000 or NovaSeq).

**Note S7. Construction of RB-TnSeq libraries in OAS795 and OAE497.**

*Variovorax* sp. OAS795 was conjugated with *E. coli* WM3064 harboring the pHLL250 *mariner* transposon vector library (strain AMD290) as explained in section 2.3, but using 50 µg/mL Km instead of 10 µg/mL Km. The final RB-TnSeq *mariner* mutant library was named *Variovorax*_OAS795_ML2. For *Rhizobium* sp. OAE497, it was conjugated with AMD290 and mutants were selected on R2A supplemented with 50 µg/mL Km, and followed by growing the library in liquid YM (yeast mannitol) supplemented with the same amount of Km. The final RB-TnSeq *mariner* mutant library was named *Rhizobium*_OAE497_ML4.

### Figures:

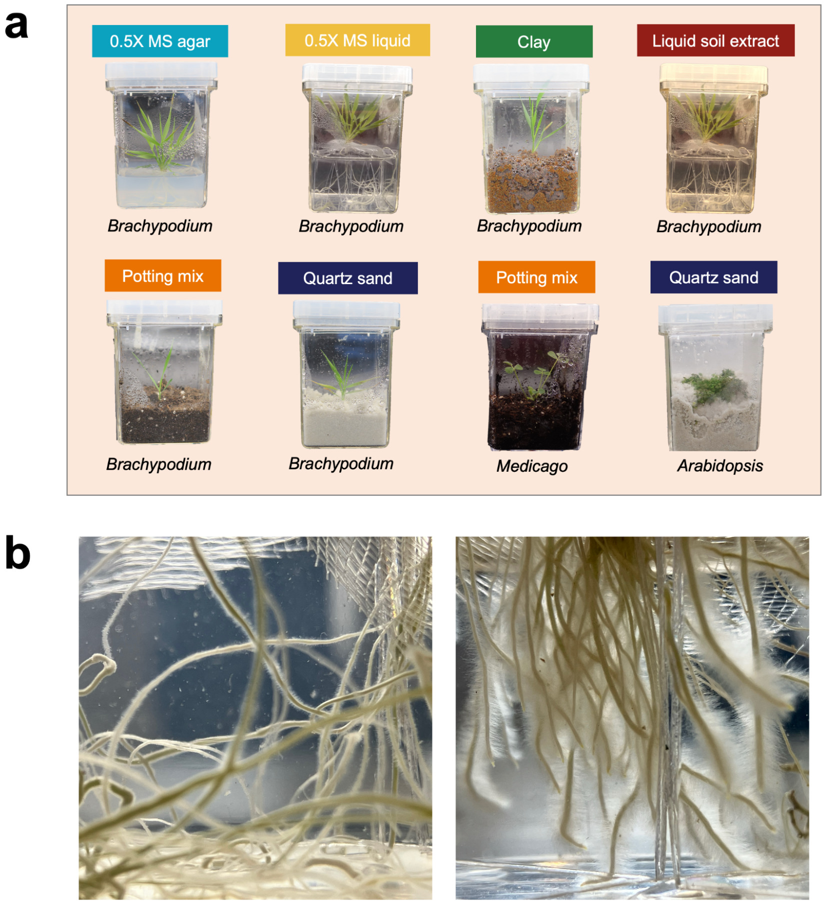


**Figure S1**. **a**) Overview of *Brachypodium distachyon* Bd21-3, *Medicago truncatula* A17 and *Arabidopsis thaliana* Col-0 plants grown in different types of substrate: 0.5X MS agar, 0.5X MS liquid, clay, liquid soil extract, potting mix, and quartz sand. **b**) Different root morphology (hair roots) observed in *Brachypodium* plants grown in 0.5X MS liquid and liquid soil extract.

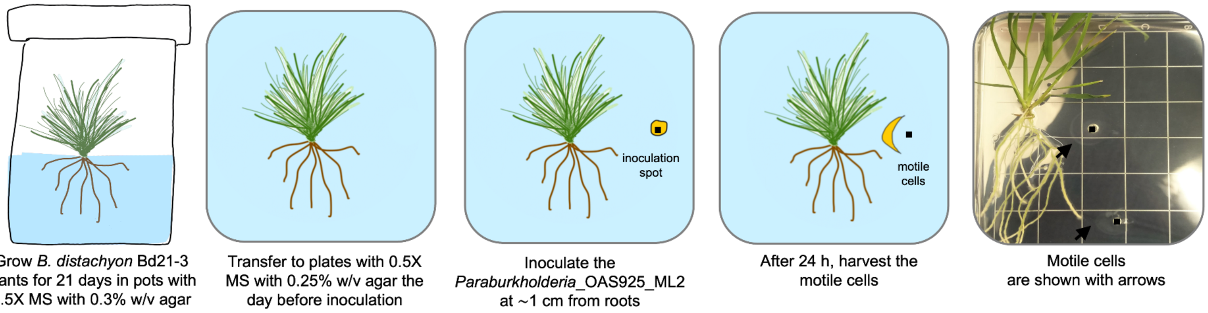


**Figure S2**. "Chemotaxis to plant root" assay. *Paraburkholderia*_OAS925_ML2 was inoculated at ~1 cm (black dot, where the bubbles appear) from 21-old-day *Brachypodium distachyon* Bd21-3 plants that were transferred from pots with 0.5X MS with 0.3% w/v agar onto plates with 0.5X MS with 0.25% w/v agar the day before inoculation. Twenty four h after inoculation, outer samples (arrows) with motile cells were harvested.

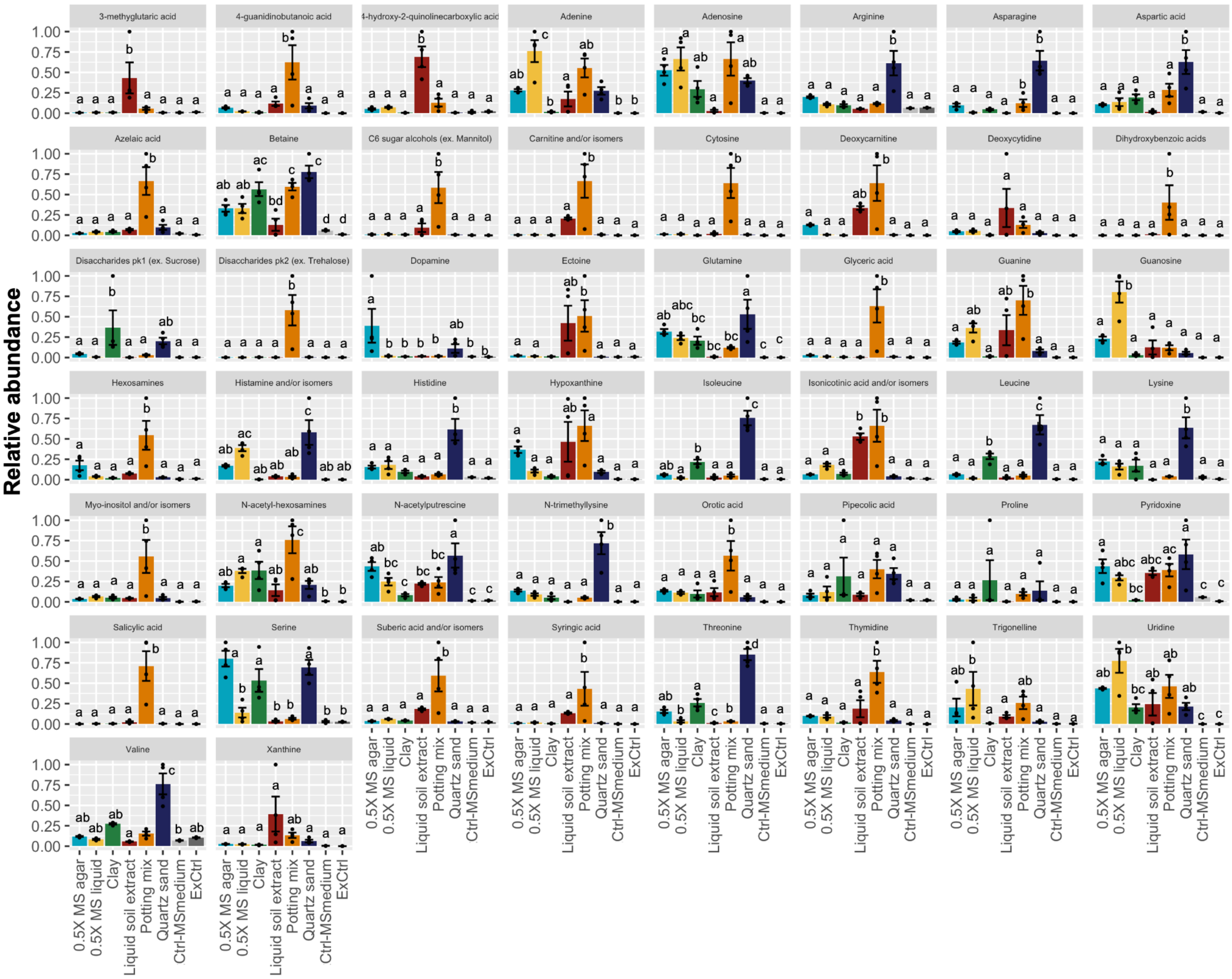


**Figure S3**. Average relative abundance of compounds across different plant growth substrates. Extraction and media controls (ExCtrl, Ctrl-MSmedium) were used to identify and remove contaminants (see Material and Methods). In some cases compound names in panels have been shortened for illustration purposes, see Table S5. Different letters indicate statistically significant differences (two-way analysis of variance (ANOVA) with Tukey's post hoc test; $P \leq 0.05$).

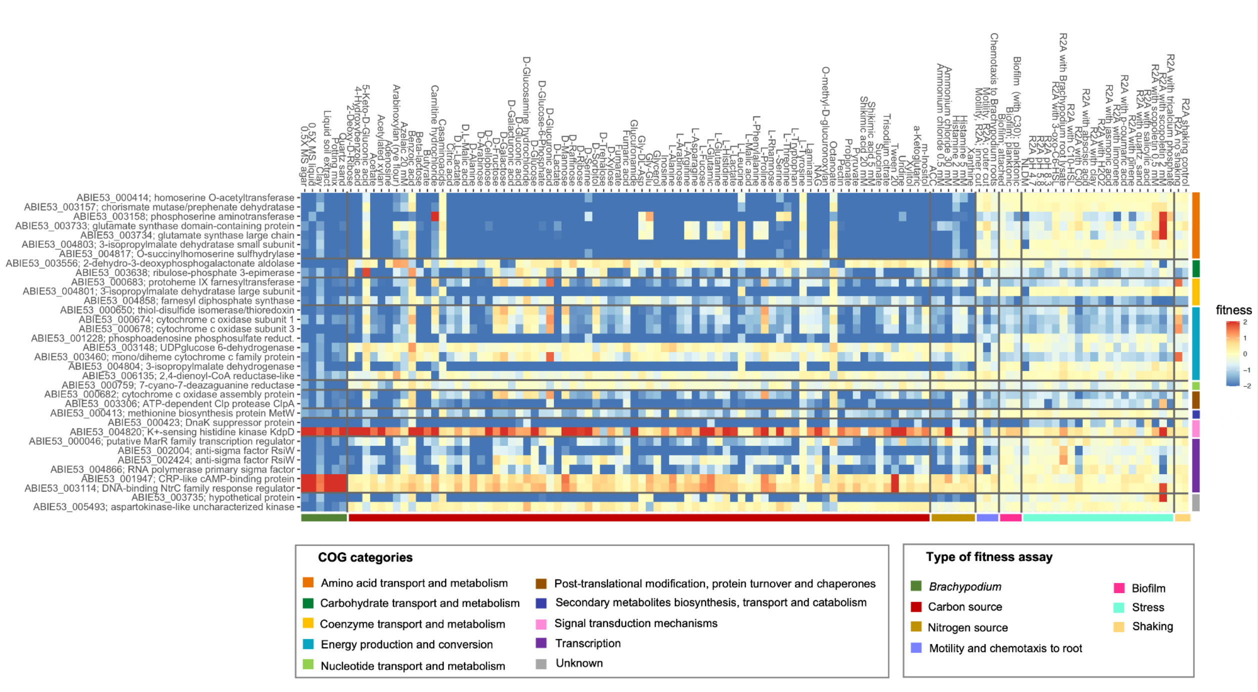


**Figure S4**. Heatmap of average fitness values for the 34 core rhizosphere colonization genes across *Brachypodium* plants grown in different plant growth substrates and different *in vitro* conditions. Gene fitness values are bounded from -2 to 2.

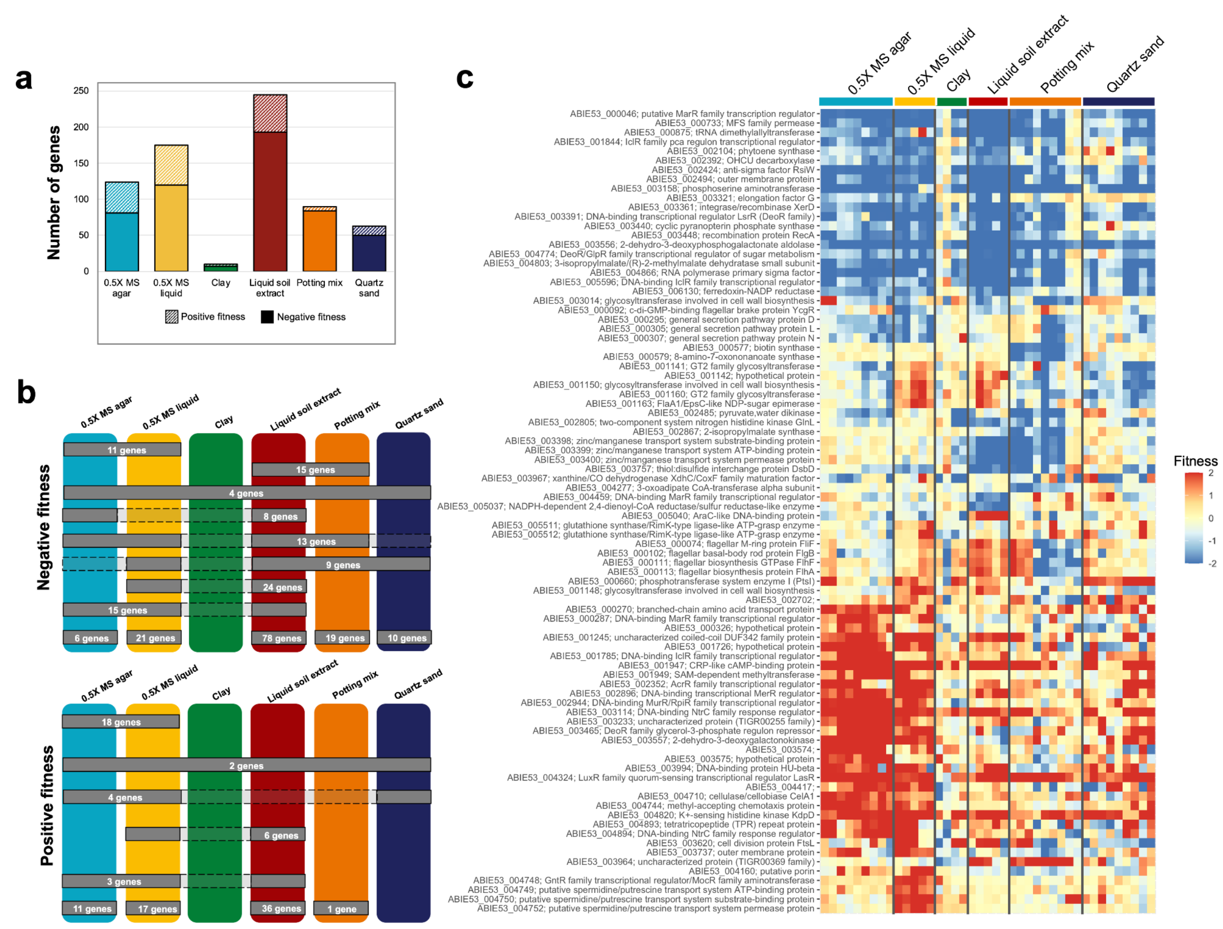


**Figure S5**. Influence of plant growth substrate on *Paraburkholderia graminis* OAS925 bacterial fitness for plant colonization. **a**) Total number of genes with negative or positive fitness values in the different substrates (|fitness| ≥ 1 and |t| ≥ 3). **b**) Shared and unique fitness colonization genes (|fitness| ≥ 1 and |t| ≥ 3) across plant growth substrates. Gray boxes show the number of shared genes across substrates; dashed empty boxes mean genes are not required for that type of substrate. For illustration purposes, numbers (n ≤ 6) corresponding to genes shared in a few substrates only are not shown in the top figure. **c**) Heatmap of fitness values across the different plant growth substrates for a subset of genes; genes with large differences in fitness between two different substrates (calculated as the |maximum fitness - minimum fitness|) are shown. Gene fitness values for each replicate experiment are plotted separately. Gene fitness values are bounded from -2 to 2.

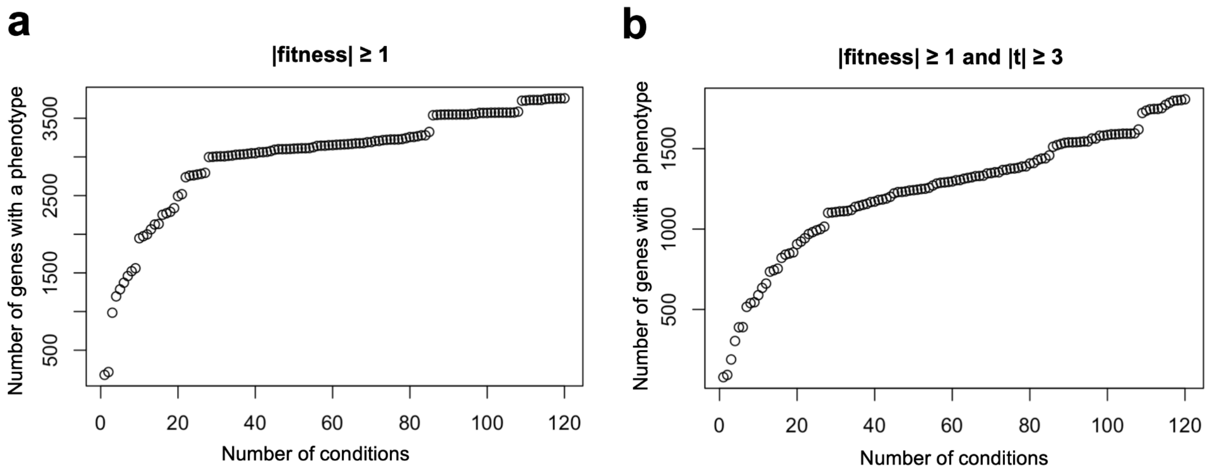


**Figure S6**. Number of genes with a fitness phenotype after conducting experiments in a given number of conditions. **a**) Criteria is |fitness| ≥ 1. **b**) Criteria is |fitness| ≥ 1 and |t| ≥ 3.

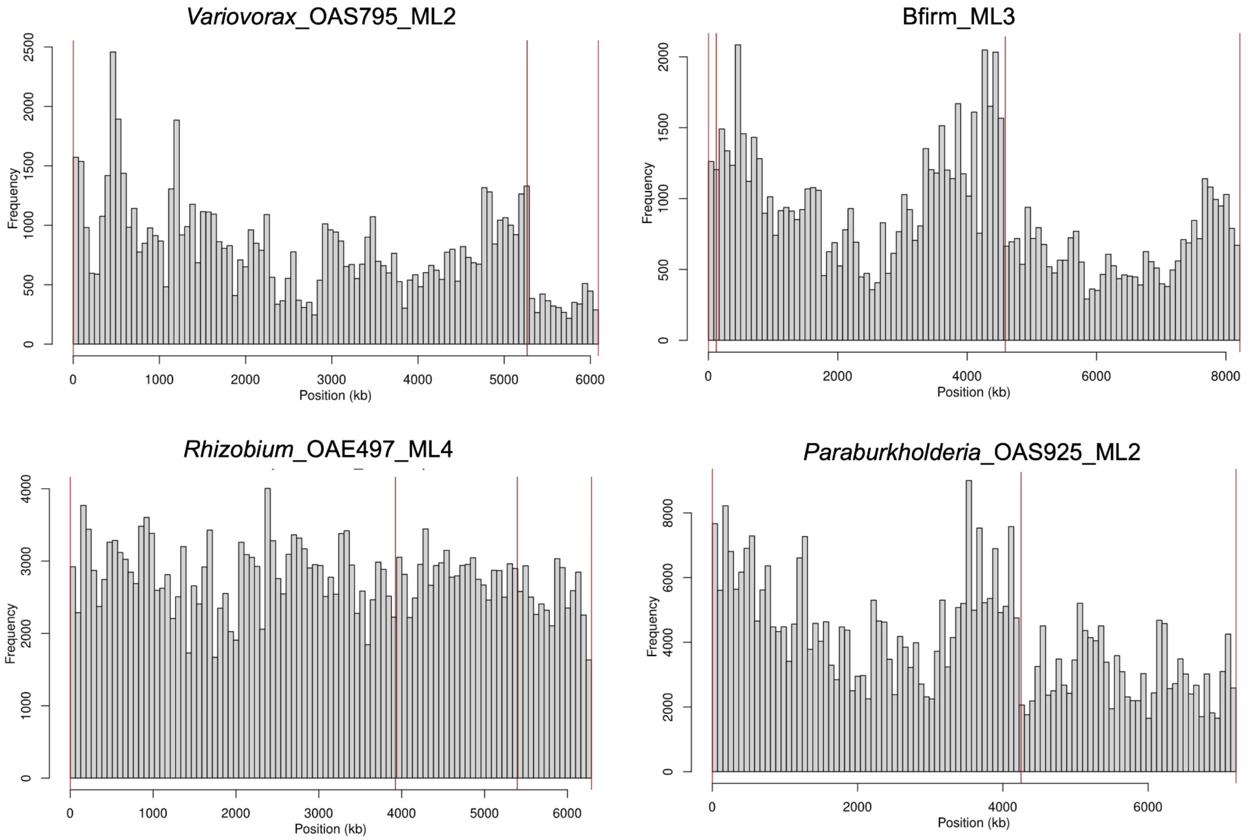


**Figure S7**. Histograms of mapped insertions for *Variovorax*_OAS795_ML2, BFirm_ML3, *Rhizobium*_OAE497_ML4 and *Paraburkholderia*_OAS925_ML2.


### Funding:

This material by m-CAFEs Microbial Community Analysis & Functional Evaluation in Soils, a Science Focus Area led by Lawrence Berkeley National Laboratory, is based upon work supported by the U.S. Department of Energy, Office of Science, Office of Biological & Environmental Research under contract number DE-AC02-05CH11231. This work used data analytical resources at the National Energy Research Scientific Computing Center, a Department of Energy Office of Science User Facility operated under contract number DE-AC02-05CH11231.
